# Coupled metalipidomics-metagenomics reveal structurally diverse sphingolipids produced by a wide variety of marine bacteria

**DOI:** 10.1101/2024.01.25.577268

**Authors:** Su Ding, F. A. Bastiaan von Meijenfeldt, Nicole J. Bale, Jaap S. Sinninghe Damsté, Laura Villanueva

## Abstract

Microbial lipids, used as taxonomic markers and physiological indicators, have mainly been studied through cultivation. However, this approach is limited due to the scarcity of cultures of environmental microbes, thereby restricting insights into the diversity of lipids and their ecological roles. Addressing this limitation, here we apply for the first time metalipidomics combined with metagenomics in the Black Sea, classifying and tentatively identifying 1,623 lipid-like species across 18 lipid classes. We discovered over 200 novel, abundant, and structurally diverse sphingolipids in euxinic waters, including unique 1-deoxysphingolipids with long-chain fatty acids and sulfur-containing groups. Genomic analysis revealed that members of 38 bacterial phyla in the Black Sea can synthesize sphingolipids, representing a fourfold increase from previously known capabilities and accounting for up to 25% of the microbial community. These sphingolipids appear to be involved in oxidative stress response and cell wall remodeling. Our findings underscore the effectiveness of multi-omics approaches in exploring microbial chemical ecology.

## MAIN TEXT

Microbial lipids are structurally highly diverse and have proven to be robust indicators of specific taxonomic groups^1^. In addition, many microbes regulate their membrane lipid composition to adapt to environmental stresses. Consequently, lipids can also be used as biomarkers for specific environmental conditions or as indicators of climate change^2–6^. The distribution of intact polar lipids have been reported in many marine settings^7–14^, lakes^15^ and soils^16, 17^. Recent developments in the field of “molecular omics”, employing non-targeted methodologies alongside computational techniques, have facilitated advancements in environmental metabolomics^18, 19^. This allows for a comprehensive characterization of all lipid molecules in the environment, the so-called metalipidome, aiding in the discovery of novel molecules^7, 20, 21^ with unknown biological sources.

Traditionally, the identification of microorganisms that synthesize a specific lipid and insights in its function have relied on microbial cultivation and further lab experiments. However, the lack of cultured representatives for the vast majority of microbes hinders a comprehensive understanding of the structural diversity of lipids and their molecular and ecological functions across the tree of life^21^. A different approach involves identifying lipids directly in the environment and inferring their biological sources and potential functions. Currently, most studies suggesting potential lipid producers in the environment rely on the co-variation of lipid abundance with molecular taxonomic markers, such as the 16S rRNA gene^22^. However, this approach can lead to erroneous conclusions, as not all microbes that co-vary with a molecular product necessarily have the capacity to produce it. Alternatively, it is possible to investigate the genetic capacity of a microorganism to synthesize a lipid compound of interest by targeting their lipid biosynthetic pathways^22, 23^. However, this method requires knowledge of the enzymes involved in the lipid biosynthetic pathways and sometimes depends on the direct detection by PCR amplification of the targeted protein-coding genes, which could induce biases^24, 25^. At present, metagenomic sequencing, combined with *de novo* assembly and genome-resolved metagenomics, provides a cultivation-independent, unbiased, and comprehensive view of the microbial community^26, 27^. In principle, this method allows for the identification of likely producers of lipids, especially if biosynthesis genes are known. However, currently, environmental metalipidomic studies have been scarce, and their integration with genomics/metagenomics is unprecedented.

In this study we present, for the first time, how combining environmental metalipidomics with metagenomics can result in the identification of novel lipids and the assignment of their microbial sources. Our study of the metalipidome of the water column of the Black Sea reveals a large structural diversity of a group of membrane lipids, i.e. sphingolipids, spanning from the surface oxygenated waters to the deep anoxic and sulfidic (euxinic) zone. By coupling this with metagenome analysis, we revealed previously largely overlooked microbial sources and suggested that the detected sphingolipids in the deep waters are associated with oxidative stress and cell wall remodeling, possibly reflecting adaptations to the euxinic environment.

## RESULTS AND DISCUSSION

The Black Sea is the largest stratified anoxic marine basin in the world, holding promise for the discovery of novel biological molecules due to its distinct microbial niches caused by redox zonation. To comprehensively characterize its complete metalipidome and its potential contributors, we collected suspended particulate matter (SPM) covering the full range of redox zones from 50 to 2,000 meters below sea level (mbsl; 15 depths): the oxic zone (50–70 mbsl), and the anoxic zone (80–2,000 mbsl), which is further subdivided in the suboxic zone (80–105 mbsl; no detectable sulfide), and the euxinic zone (110– 2,000 mbsl; sulfidic)^28, 29^. Prior investigations^28^ of the physicochemical characteristics of the Black Sea water column (see Supplementary Fig. 1) revealed that oxygen concentrations exhibited a noticeable decline from the surface layer downward, decreasing from ∼78 µM at 50 mbsl to levels under the detection limit around 80 mbsl. The same trend was also found for nitrate, which had a concentration of ∼1.3 µM at 50 mbsl and concentrations under the detection limit at ∼90 mbsl. Sulfide was not detected in the surface layer and appeared at ∼110 mbsl (0.85 µM), which marks the upper boundary of the euxinic zone (here defined as sulfide concentration > 0.2 μM). Lipids and DNA were extracted based on previous methods^8, 28^.

### Characterization of the metalipidome of the Black Sea

Lipid extracts obtained from the SPM across the water column of the Black Sea were analyzed with Ultra High-Pressure Liquid Chromatography (UHPLC) coupled with high-resolution mass spectrometry (HRMS)^8^. The selected analytical window for the MS/MS mass spectra of ion components detection was *m/z* 350–2,000, targeting the relatively large components of the environmental metabolome. To characterize the chemical diversity of this metalipidome, the MS/MS mass spectra of ion components were subjected to molecular networking based on their cosine similarities (Fig. 1a). A clustering of ion components in the molecular network suggests similar molecular structures, for example as analogs with an identical polar headgroup or similar core that differ through simple transformations such as alkylation, unsaturation, and glycosylation. Although only a small fraction (less than 2%) of the components in the metalipidome could be annotated through reference database searches, we successfully classified or putatively identified 1,623 lipid species into 18 major lipid classes within the molecular network. This was based on manual inspection of specific abundant components in the subnetworks^7^ (Fig. 1b).

**Fig. 1.**
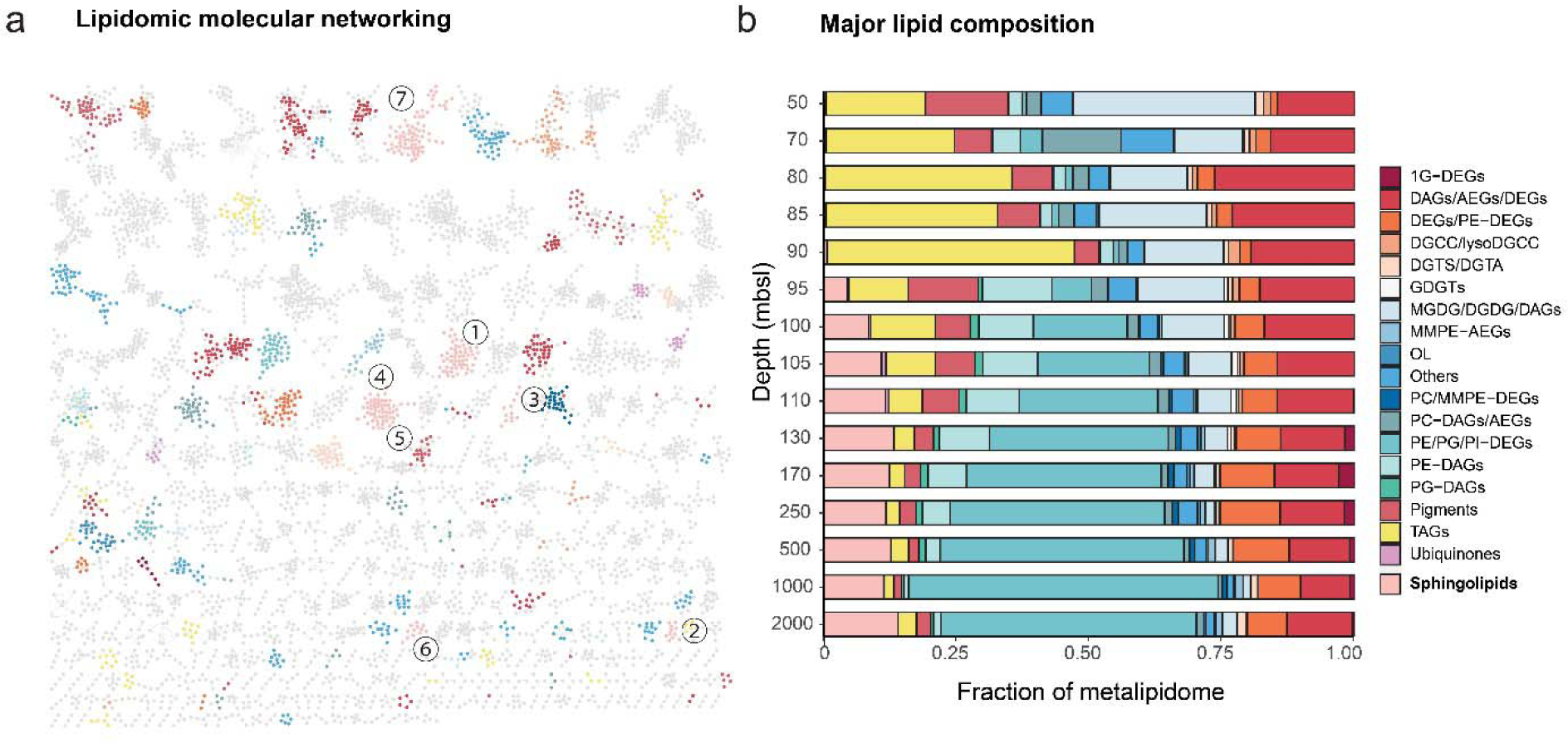
Metalipidome composition of the Black Sea. (**a**) Molecular network of the metalipidome of the water column (combining all SPM samples from different water depths). Nodes represent MS/MS mass spectra of ion components within an analytical window of *m/z* 350–2,000, which are connected based on spectral similarity (cosine > 0.6). Nodes with colors are major lipid classes shown in **b** and numbers refer to the seven sphingolipid classes shown in Fig. 2d. The spatial orientation of the nodes in the MS/MS network is randomly generated by Cytoscape^42^ and does not relate to relationships between the subnetworks. (**b**) The relative abundance of major lipid classes in the water column of the Black Sea. Relative abundances of lipids were calibrated using the response factor of external standards (see ‘experimental procedure’ for details). Major lipid classes that did not have a relative abundance >1% in multiple SPM samples are classified as ‘other’. Lipid abbreviations: 1G-DEGs, monoglycosyldietherglycerols; DAGs, diacylglycerols; DEGs, dietherglycerols; AEGs, acyletherglycerols; TAGs, triacylglycerol; DGTS, diacylglycerylhydroxymethyltrimethyl-(N,N,N)-homoserine; DGCC, diacylglycerylcarboxyhydroxymethylcholine; DGTA, diacylglyceryl-hydroxymethyl)-tri-methyl-b-alanine; OL, ornithine lipid; PC, phosphatidylcholine; PE, phosphatidylethanolamine; PG, phosphatidylglycerol; PI, phosphatidylinositol; MMPE, phosphatidyl-(N)-methylethanolamine; MGDG, monoglycosyldiacylglycerol; DGDG, diglycosyldiacylglycerol; GDGT, glycerol dialkyl glycerol tetraethers.

The 18 major classes of lipids (Fig. 1b; abbreviations in the caption of Fig. 1) included core lipids (DAGs, AEGs, and DEGs), phospholipids (PE, PG, PI and MMPE headgroups connected to DAGs, AEGs, and DEGs), glycolipids (1G-DEGs, MGDG, DGDG), GDGTs, betaine lipids (DGCC and lysoDGCC), DGTS/DGTA, ornithine lipids (OLs), pigments, triacylglycerols (TAGs), ubiquinones, and sphingolipids. The lipids with the highest summed relative abundance at 50–90 mbsl depths were TAGs, constituting 19-46% of all lipids, following by glycolipids (13-34%), and DAGs (15-26%) (Fig. 1b). TAGs are energy storage molecules of eukaryotic algae such as *Euglenophyceae*, *Cryptophyceae*, and *Eustigmatophyceae*^30^, and their relative abundance falls below 5% in the euxinic zone.

Unexpectedly, sphingolipids reached an unprecedented high abundance of up to 14% of the metalipidome in the deep euxinic waters (Fig. 2b). Sphingolipids are composed of a sphingosine backbone linked to a fatty acid via an amide bond (Fig. 2). They play essential roles in eukaryotes as fundamental building blocks of the cell membrane and serve vital functions in cellular signaling and organization of lipid rafts^31–33^. However, even though sphingolipids in marine environments were previously thought to be primarily produced by eukaryotic algae and their associated viruses^34–36^, they accounted for < 0.5% in the oxic and upper suboxic zone (50–90 mbsl), the primary niches for eukaryotes in the Black Sea. In contrast, sphingolipids had a mean relative abundance of 12% in the euxinic zone, making them the third most abundant lipid class in these deep waters, after phospholipids (mainly PE/PG/PI-DEGs; 8–58%) and core lipids (mainly AEGs and DEGs; 9–18%), strongly suggesting a prokaryotic source. However, sphingolipids are thought to be rare in bacteria, although recently their presence has been inferred in 9 bacterial taxonomic phyla^37^ (further information about bacterial sphingolipids can be found in the Supplementary Note). A previous study identified a single subclass of sphingolipids in the Black Sea, glycosylceramides (Gly-Cers), and suggested that they were produced by (facultative) anaerobic bacteria because of the high abundance of these lipids in the anoxic deep waters^12^. Here, we identify, by using a combined metalipidomics-metagenomics approach, the complexity in structural diversity and distribution of the sphingolipids and unravel their microbial sources.

**Fig. 2.**
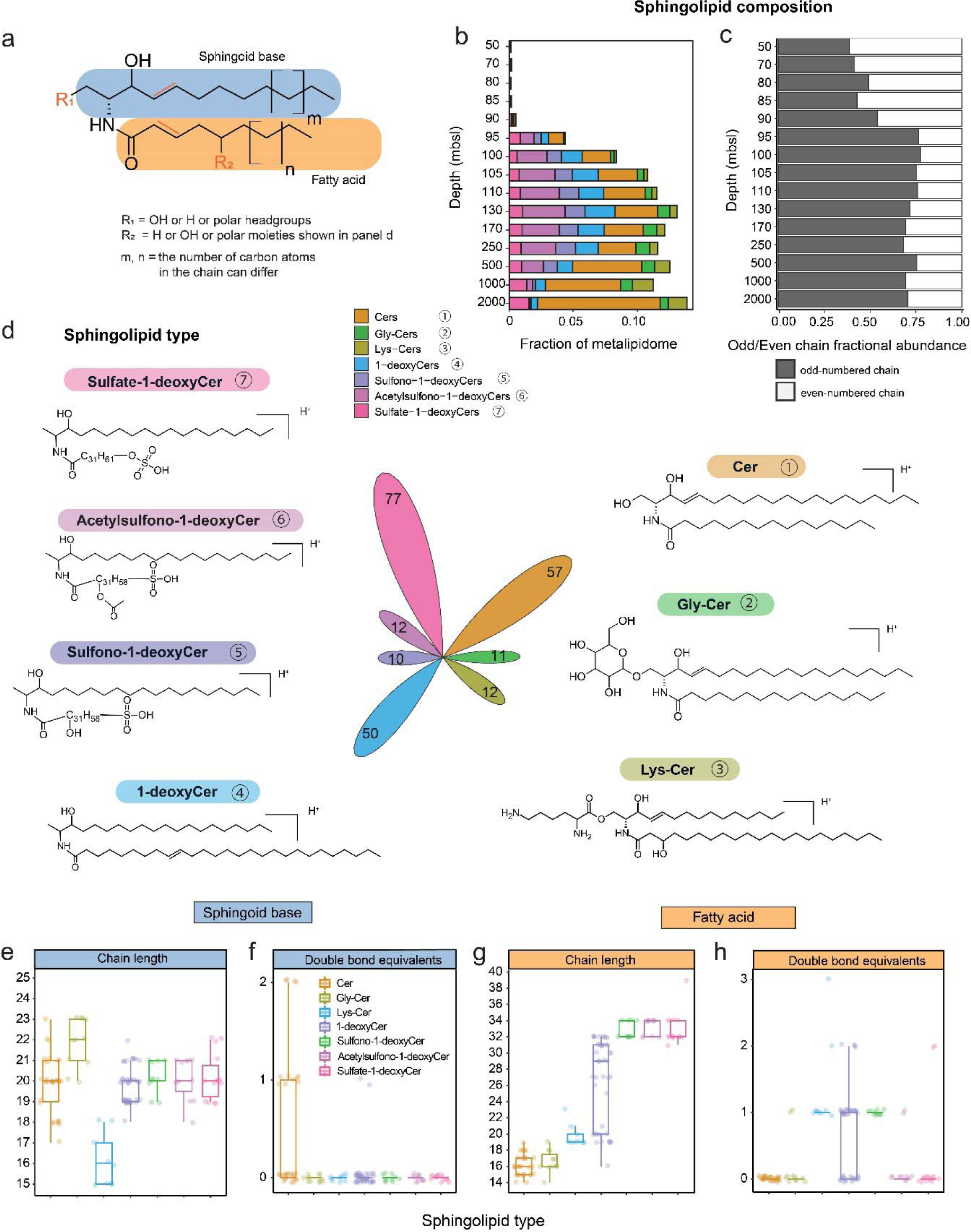
Sphingolipids in the Black Sea water column. (**a**) General structure of sphingolipids consisting of a sphingoid base linked to a fatty acid via an amide-bond. The number of carbon atoms of the sphingoid base and fatty acid moieties ranged from 15 - 22 and 14 - 34, respectively. (**b**) The relative abundance of seven sphingolipid classes identified in the metalipidome. (**c**) The variation in the fractional abundance of odd- and even-numbered chain sphingolipids with depth in the water column of the Black Sea. An ‘odd-numbered chain’ sphingolipid contains either an odd chain sphingoid base and/or an odd chain fatty acid. An ‘even-numbered chain’ sphingolipid contains even chains of both the sphingoid base and fatty acid. (**d**) Structures of each representative sphingolipid class. The flower plot area with the numbers in the center represents the number of distinct ion components within each subclass. The ranges of (**e**) chain length and (**f**) the number of double bond equivalents (DBEs) of the sphingoid base for the various sphingolipid classes. The ranges of (**g**) chain length and (**h**) the DBEs of the fatty acid base. Out of the 229 identified sphingolipids, we were able to assign molecular structures with confident information on chain-length carbon atoms and DBEs to 122 (see Source dataset). These 122 sphingolipids were subjected to detailed Fig. 2e-h analysis. In panel a, for the class of Cer, R_1_ = OH, R_2_ = H; for the class of Gly-Cer, R_1_ = Gly, R_2_ = H; for the class of Lys-Cer, R_1_ = Lys, R_2_ = OH; for the class of 1-deoxyCer, R_1_ = H, R_2_ = H; for the class of Sulfono-1-deoxyCer, R_1_ = H, R_2_ = SO_3_H and OH (the position of these two groups cannot be determined based on the mass spectra and they are not connected to each other); for the class of Acetylsulfono-1-deoxyCer, R_1_ = H, R_2_ = SO_3_H and CH_3_COO (the position of these two groups cannot be determined based on the mass spectra and they are not connected to each other); for the class of Sulfate-1-deoxyCer, R1 = H, R2 = SO_4_H.

### Structural diversity and distribution of sphingolipids

The metalipidomic approach revealed that the Black Sea contained 229 distinct structures of sphingolipids in the deeper suboxic waters (Fig. 2). These sphingolipid structures varied in polar headgroups/moieties, chain length, and degree of unsaturation. The 229 distinct sphingolipids structures can be divided into 7 subclasses: ceramides (Cers; also including saturated ceramides), Gly-Cers, lysine-ceramides (Lys-Cers), 1-deoxyceramides (1-deoxyCers), sulfono-1-deoxyceramides (sulfono-1-deoxyCers), Acetylsulfono-1-deoxyceramides (Acsulfono-1-deoxyCers) and sulfate-1-deoxyceramides (sulfate-1-deoxyCers) (Fig. 2d). We also examined the core lipid composition of each subclass of sphingolipids, focusing on the number of carbon atoms and double bond equivalents (DBEs) in their sphingoid base and their fatty acid moiety (Fig. 2 and Supplementary Fig. 2), as these two parameters represent a large fraction of the structural diversity within the molecular subnetworks.

The Cers subnetwork comprised 57 different structures, and it represented the most abundant sphingolipid subclass in the euxinic zone, accounting for 3-10% of total metalipidome. Approximately half of the Cer structures identified (median 46%) exhibited one DBE in their sphingoid base, indicating that the remaining Cers were dihydroCers (Fig. 2c). Gly-Cers had their highest summed relative abundance at 130-1,000 mbsl, and the core lipid composition of Gly-Cers was largely consistent with previously identified sphingolipids^12, 14, 38^, with sphingoid bases ranging from 19 - 23 carbon atoms and C_14_ - C_19_ acyl moieties (Fig. 2e,g). Lys-Cers have not been reported before in natural environments, and had their highest summed relative abundance in the deepest waters. The novel Lys-Cers had the shortest base sphingoid chain length (median C_17_; Fig. 2e).

The four groups of 1-deoxyCers contained 149 structures, including many that have not been previously reported in the natural environment nor in microbial cultures (see below). Together they contributed 2-8% of total lipids in the euxinic zone. 1-deoxyCers lack the C1-hydroxyl group of the classical ceramides on their sphingoid backbone. Due to the lack of the hydroxy moiety, these uncommon ‘headless’ sphingolipids were hypothesized to be unable to form complex sphingolipids^39, 40^ (see Gly-Cers and Lys-Cers in Fig. 2d and the polar functional groups at their C1-OH position). Our identification of the other three 1-deoxyCer subclasses shows, however, that they are in fact capable of bearing polar moieties, such as sulfate, sulfono and acetylsulfono on their fatty acid chains. The relative abundance of all the 1-deoxyCers-associated sphingolipids increased to a maximum at 130 mbsl and decreased again with depth, except for sulfate-1-deoxyCers, which had their highest summed relative abundance in the deepest waters. The four subclasses of 1-deoxyCer-associated sphingolipids exhibited sphingoid bases that were similar to each other, with 18 to 22 carbon atoms and mostly no unsaturations (Fig. 2e,f). In contrast, 1-deoxyCers contained the widest range in fatty acid chain lengths of all sphingolipid subclasses with 16 to 32 carbon atoms, while the three subclasses with additional polar moieties had similar fatty acid chain lengths consisting mostly of 32 to 34 carbon atoms (Fig. 2g). The longest fatty acid chain observed in the Black Sea was a C_39_ from a Sulfate-1-deoxyCer. 1-deoxyCers had between 0 and 2 DBEs on their fatty acid chains, all Sulfono-1-deoxyCers possessed 1 fatty acid chain DBE, whereas acsulfono-1-deoxyCers and sulfate-1-deoxyCers predominantly lacked fatty acid chain DBEs.

The sphingoid bases of 1-deoxyCers (1-deoxysphinganines) were initially discovered in the marine bivalve *Mactromeris polynyma*^41^, and have been associated with other eukaryotes such as fungi, other bivalves, sponges, and mammals^39, 40, 42, 43^. The very long (i.e., >31) fatty acid chains of the 1-deoxyCer-associated sphingolipids in the Black Sea have to our knowledge not been reported from any sphingolipid in environmental microbiomes. Sphingolipids containing ‘long-chain fatty acids’ (LCFAs) of eukaryotic origin with a chain length of 22 carbon atoms have been found in mammals^44–49^, plants^50–54^, invertebrates^55^ and fungi^53, 56–58^. These eukaryotic LCFAs consistently exhibit an even number of carbon atoms (Supplementary Fig. 3). In marine environments, the eukaryotic alga *E. huxleyi* and its associated viruses have been observed to synthesize Gly-Cers containing LCFAs with 22 carbon atoms^34, 35, 59^. The 1-deoxyCer-associated sphingolipids of the Black Sea contain fatty acids with chain lengths up to 39 carbon atoms. Over 50% of these LCFAs possessed an odd chain length (Fig. 2c and Supplementary Fig. 3). The absence of odd-chain LCFAs in known sphingolipids of eukaryotes, and the observation that odd-chain fatty acids in intact polar lipids occur predominantly in bacteria^16, 60^, moreover suggests that the sphingolipids in the deep waters of the Black Sea are produced by bacteria.

### The Black Sea harbors a diverse bacterial community of sphingolipid producers

To comprehensively identify likely microbial producers of the unusual sphingolipids in the Black Sea, we deeply sequenced the metagenomes of the SPM that was used for lipid analysis, followed by de novo assembly and binning of the assembled scaffolds in metagenome assembled genomes (MAGs). Next, we queried the predicted protein sequences on the ∼1.64 x 10^8^ assembled scaffolds for the key Cer biosynthesis enzyme serine palmitoyltransferase (Spt). Spt catalyzes the condensation of L-serine and palmitoyl-coenzyme A as the first step in the biosynthesis of the ceramide backbone, and can alternatively use alanine as a substrate instead of serine in the biosynthesis of the 1-deoxyCer backbone (Supplementary Fig. 4)^37, 61–64^. We hypothesize that Spt can also use other amino acids as a substrate, like lysine to generate Lys-Cer. Spt is part of the alpha-oxoamine synthases protein family that contains diverse enzymatic activities^6^ and even with our stringent search criteria (*e*-value ≤ 1 x 10^-70^, query coverage ≥ 70%) not all Spt homologs (Supplementary Data 1) may have Spt enzymatic activity. To identify homologs with likely Spt activity, we constructed a phylogenetic tree of the 8,524 Black Sea hits and known alpha-oxoamine synthases (Fig. 3a; Supplementary Data 2), hypothesizing that phylogenetic clustering with known Spt sequences implies conserved enzymatic activity. We also queried the predicted protein sequences for bacterial ceramide synthase (bCerS) and ceramide reductase (CerR), that together with Spt constitute the known bacterial (1-deoxy)Cer backbone biosynthesis pathway (Supplementary Fig. 4) and are often found clustered on the genome^37, 64^. By using these search criteria for sphingolipid biosynthesis genes, we aim to more accurately pinpoint potential producers based on their genomic capability.

**Fig. 3.**
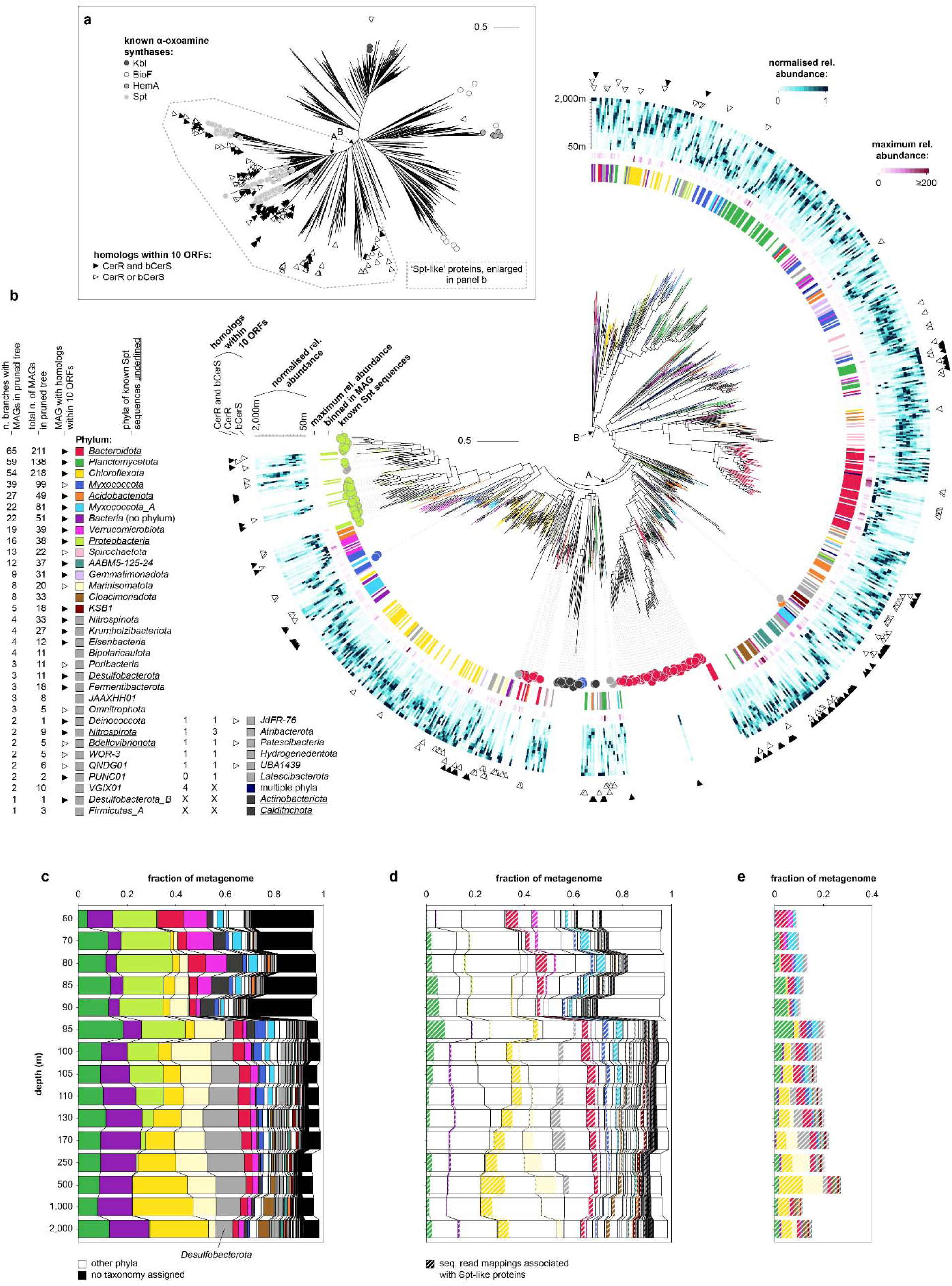
Potential sphingolipid producers in the Black Sea based on the presence of Spt-like proteins encoded on metagenomes. (**a**) Phylogenetic tree of Black Sea Spt homologs and reference sequences. The tree contains 2,666 representative protein sequences that represent 8,524 hits from the Black Sea, 273 known Spt sequences, and 16 known homologous sequences with different enzymatic activities. Co-localization of genes encoding CerR and bCers with Spt homologs on the scaffold is indicated with triangles. The clades containing all known Spt sequences (clade A) and most sequences with CerR and bCerS encoded in close proximity on the scaffold (clade B; ‘Spt-like’ proteins) have high bootstrap support (≥ 98). An annotated tree with bootstrap support can be found in Supplementary Data 2. (**b**) Zoom in on clade B. Taxonomic annotation at the rank phylum of medium to high quality MAGs (> 50% completeness, < 10% contamination) in which Spt-like proteins are binned (1,271 in total) is indicated with color-coding of the branches and around the tree, Spt-like proteins encoded on unbinned scaffolds are not colored. Taxonomic affiliation of known Spt sequences is indicated in circles around the tree. Scale bars in panels a and b show the mean number of substitutions per site. (**c**) Community profile based on sequencing read mapping to scaffolds. Taxonomic annotation of the MAG is used if the scaffold is binned, and of the scaffold otherwise. Stacked bars add up to less than 1 because not all reads can be mapped to scaffolds. (**d**) Idem to panel c but with the fraction of sequencing read mappings colored that are associated with a Spt-like protein via the MAG or unbinned scaffold. (**e**) Idem to panel d but stacked. Spt, serine palmitoyltransferase; bCerS, bacterial ceramide synthase; CerR, ceramide reductase; ORF, open reading frame; n., number; MAG, metagenome assembled genome; rel., relative; tax., taxonomy; seq., sequencing. Source data for the figure can be found in Supplementary Data 2 and 3.

The phylogenetic tree of Spt homologs contained a clade that included all previously known Spt sequences (Fig. 3a,b, clade A) and a larger clade that included proteins with CerR and bCerS homologs encoded in close proximity on the same scaffold (Fig. 3a,b, clade B). We considered this larger clade to potentially code for an Spt-like enzymatic activity, and the microorganisms that encode these proteins (called ‘Spt-like’ proteins from hereon) to have biosynthetic potential to produce sphingolipids. Our phylogenetic approach to identifying the enzymatic activity of Spt homologs showed many cases in which the gene encoding the Spt-like protein was not co-localized on the genome with genes encoding bCerS and CerR. For example, none of the 33 *Cloacimonadota* MAGs had bCerS or CerR encoded within a distance of 10 open reading frames (ORFs) of the gene encoding the Spt-like protein (Fig. 3b), possibly indicating rearrangement of the sphingolipid biosynthesis genomic cluster, or replacement of the enzymatic activity of bCerS and CerR by alternative non-homologous proteins. It has previously been observed that the three genes are also not encoded in close proximity on the genomes of sphingolipid-producing members of *Bacteroidota*^37^, indicating that genomic co-localization is not required for sphingolipid biosynthesis. Alternatively, fragmented assembly and incomplete binning may result in an underestimation of the genomic co-localization of these genes.

The clade with Spt-like proteins encompasses a much broader phylogenetic diversity of Spt than previously identified, supporting a diverse community of potential sphingolipid producers present throughout the Black Sea water column (Fig. 3b). Of the 5,069 medium to high-quality MAGs (>50% completeness, <10% contamination^65^) obtained, 1,271 encoded Spt-like proteins as defined above. These MAGs have diverse taxonomic affiliations encompassing 38 bacterial phyla, expanding the known diversity of potential sphingolipid-producing phyla from 9 (identified in ref. ^37^ based on genomic presence of Spt, bCerS, and CerR homologs; see Supplementary Data 2 for the taxonomic annotation of these sequences based on the Genome Taxonomy Database) to 40 (Fig. 3b). No archaeal MAGs were identified encoding Spt-like proteins, suggesting that archaea in the Black Sea do not produce sphingolipids or use a different biosynthetic pathway. Whereas we cannot exclude the possibility that some Spt-like sequences were erroneously binned and thus incorrectly associated with a phylum, the high number of MAGs encoding Spt-like proteins for many phyla (e.g., 138 *Planctomycetota*, 218 *Chloroflexota*, 22 *Verrucomicrobiota*, 33 *Cloacimonadota*; Fig. 3b) suggest that this is unlikely to influence our observation that the Black Sea harbors a diverse and previously unknown community of sphingolipid producers.

### Sphingolipid producers are abundant and have distinct niches

Because we targeted all assembled scaffolds of the deeply sequenced metagenomes, our homology searches provide a comprehensive overview of the potential sphingolipid producers in the water column, with only microorganisms with low relative abundance, whose DNA was not sequenced or assembled into scaffolds, overlooked. We generated taxonomic abundance profiles based on mapping of the sequencing reads to the assembled scaffolds, inheriting the taxonomic annotation of the MAG when possible or else of the assembled scaffold, an approach that has been shown to give an accurate and comprehensive view of the microbiome^66^ (Fig. 3c). We similarly generated profiles for the mappings that could be associated with Spt-like proteins via the MAG or unbinned scaffold (Fig. 3d,e). This approach revealed a high abundance of potential sphingolipid producers based on the presence of the Spt-like proteins throughout the water column, with a minimum of 7% of the total sequenced DNA of the microbial community in the oxic zone and upper suboxic zone attributable to potential sphingolipid producers, and up to 25% in the euxinic zone (Fig. 3e).

Even though we found potential sphingolipid producers in many phyla, in most cases not all of their members had the potential to synthesize sphingolipids. For example, the 38 *Proteobacteria* MAGs that are potential sphingolipid producers (Fig. 3a) represented only a minority of total sequenced *Proteobacteria* DNA (Fig. 3d). In contrast, a large fraction of *Bacteroidota* sequenced DNA which was abundant throughout the water column and of *Myxococcota_A* sequenced DNA which was abundant in the oxic and suboxic zone was attributed to potential sphingolipid producers (Fig. 3d). The abundance profiles of other phyla showed niche-based differentiation of individual members. For example, even though *Planctomycetota* were abundant throughout the water column, potential sphingolipid-producing members of the phylum peaked in the redoxcline. Potential sphingolipid-producing members of *Desulfobacterota* peaked in the upper euxinic zone, and of *Chloroflexota* and *Marinosomatota* in the deep waters (Fig. 3d). These microbial abundance patterns likely represent specific niches in the stratified Black Sea in which sphingolipid producers reside, as ultimately reflected in the abundance profiles of the different sphingolipid subclasses.

Finally, we addressed the possibility that (some of) the unusual sphingolipids were produced by eukaryotes. None of the scaffolds and MAGs that contained any Spt homolog, including the sequences that fell outside the selected clade of likely Spt enzymatic activity, had a eukaryotic taxonomic annotation (Supplementary Data 1). In addition, we queried the predicted protein sequences with known eukaryotic Spt sequences. This homology search revealed 26 hits that were not identified with our original bacterial Spt homology searches (Supplementary Data 1). The hits had the highest relative abundance in the top water layer and a low relative abundance overall (Supplementary Fig. 5c), and are thus unlikely to have significantly contributed to the observed lipid profiles in the suboxic and euxinic waters. We conclude that the sphingolipids in the Black Sea are most likely produced by the bacterial community, which showed niche-based differentiation in the stratified water column.

### Identification of potential producers of distinct sphingolipids based on co-abundance

In order to identify potential producers of distinct sphingolipids, we correlated the relative abundance profiles of the 229 sphingolipid ion components with the relative abundance profiles of the 1,455 scaffolds that encode Spt-like proteins and that were binned in MAGs (Fig. 4). 133 sphingolipid structures had a Spearman rank correlation coefficient ≥ 0.9 with 728 binned scaffolds and were visualized in a network (Fig. 4d). Connections between the most abundant lipids (Supplementary Fig. 5a) and most abundant MAGs (Supplementary Fig. 5b), revealed that 1-deoxyCers, sulfono-1-deoxyCers, and acsulfono-1-deoxyCers, which were dominant in the upper euxinic zone (110–250 mbsl), may be primarily produced by abundant *Desulfobacterota* and/or *Bacteroidota* (Fig. 4). *Bacteroidota* are known to synthesize sphingolipids in the mammalian gastrointestinal tract which modulate the host’s immune response^63^, where they have been observed to synthesize 1-deoxyCers. *Desulfobacterota* is a phylum of sulfate-reducing bacteria and had the highest relative abundance of any microorganism that encodes Spt-like proteins 100–130 mbsl (Supplementary Fig. 5b).

**Fig. 4.**
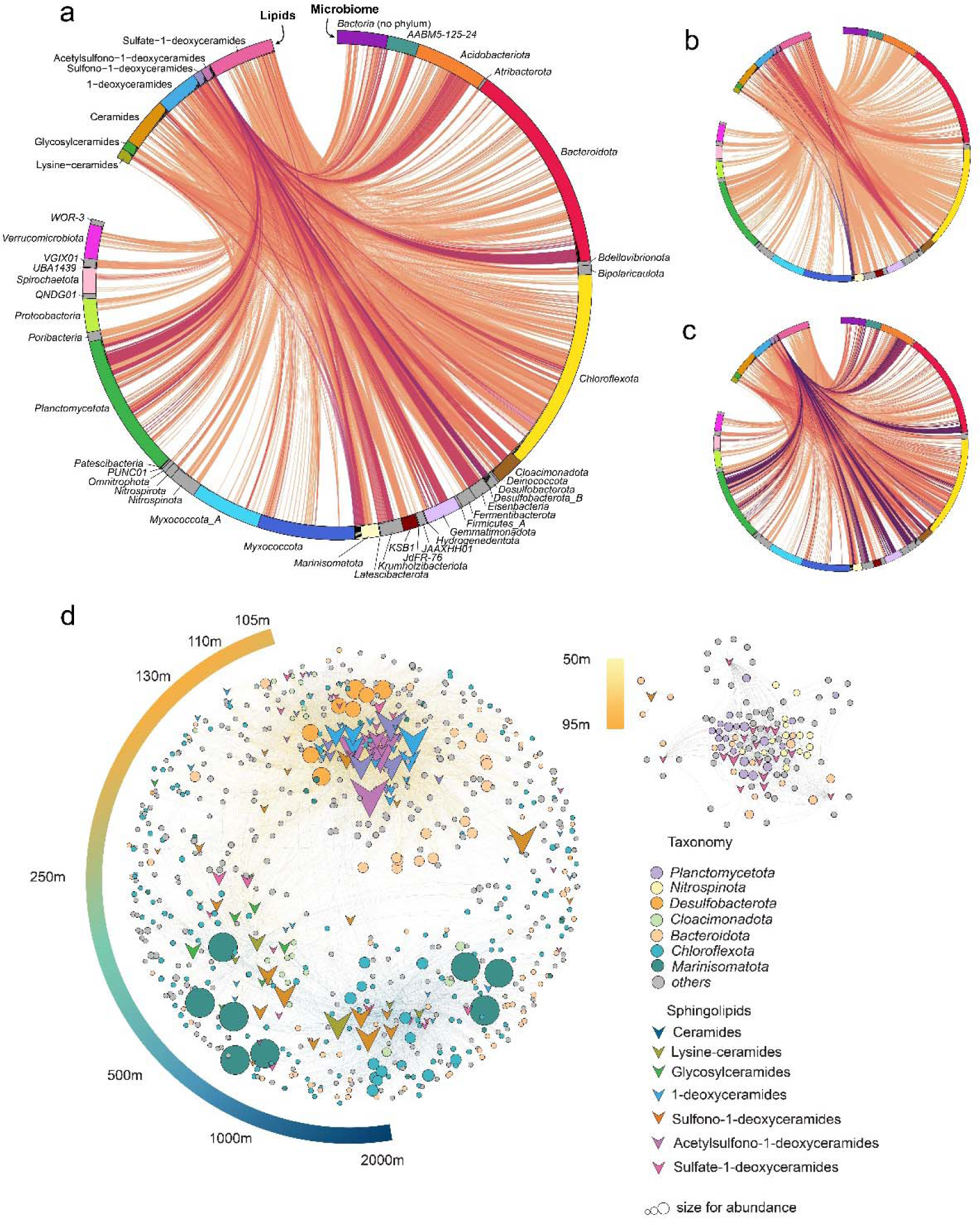
Correlations between the abundance profile of 229 sphingolipids and 1,455 scaffolds that contain Spt with likely Spt activity and that were binned in MAGs. (**a**) Lines depict Spearman rank correlations coefficients ≥0.9 between the abundance profile of the lipid and the relative abundance profile of the scaffold. The line color represents the relative abundance of the scaffolds and of the lipids, where darker colors show connections between both high abundant lipids (summed across the water column) and high abundant scaffolds (summed across the water column). Colors scale between 0 (orange; a summed relative abundance of 0 for both lipids and scaffolds) and 2 (purple; a summed relative abundance of 1 for both lipids and scaffolds). (**b**) Idem to panel a but with the lines colored according to scaffold abundance, where darker colors show connections with scaffolds that have a high summed abundance across the water column. (**c**) Idem to panel a but with lines colored according to lipid abundance, where darker colors show connections with lipids that have a high summed abundance across the water column. (**d**) Network visualization of the same data. Edges with a Spearman correlation coefficient > 0.9 are shown. The left stripe with color gradient stands for the sphingolipid abundance weight across the water column of the Black Sea (details can be found in Online Methods). The abundance weight ( ) is determined as a factor of the dominant distribution of a specific sphingolipid or scaffold across the water column. For instance, with a value close to 9 represents this sphingolipid or scaffold that has a dominant abundance at the upper euxinic zone (∼110 mbsl), such instances are denoted by the color orange on the gradient stripe. Conversely, when approaches a value of 15, it indicates a predominant distribution in the deep waters (∼2000 mbsl) and is consequently represented b the color dark blue on the stripe.

Cers, Gly-Cers, and Lys-Cers had their highest relative abundance in the deep waters (500–2,000 mbsl). These lipid subclasses showed strong correlations with *Marinisomatota* and *Chloroflexota* (of the taxonomic class *Anaerolineae*), which occurred abundantly at these depths (Fig. 4, Supplementary Fig. 5b). *Marinisomatota* is a highly diverse bacterial phylum predicted to be involved in sulfur and nitrogen cycles^67–69^, and had the highest relative abundance of any microorganism that encoded an Spt-like protein across the water column, with a maximum at 500 mbsl. *Anaerolineae* have been found in marine sediments as chemoheterotrophs, where they may use sulfur, iron, or manganese as electron sources for growth and survival^70–72^. Both *Marinisomatota* and *Chloroflexota* have not previously been identified as potential sphingolipid producers. Finally, sulfate-1-deoxyCers showed connections with a variety of bacteria spanning the upper euxinic zone to the bottom waters (Fig. 4), including the aforementioned phyla, suggesting that they may be produced by multiple anaerobic bacterial species in multiple niches.

### Potential role of sphingolipids in the Black Sea

Next, to assess the potential role of sphingolipids in the environment, we used the 5,069 medium-to high-quality MAGs to identify genomic functions that were positively or negatively associated with potential sphingolipid producers throughout the 15 depths in the water column (Online Methods). Positively associated (enriched) functions may be involved in sphingolipid biosynthesis, or be related to an ecological role of sphingolipids in the environment. Negatively associated (depleted) functions may point to a function that the sphingolipids replace. We focused on functions that were shared by a large fraction of the phyla that were potential sphingolipid producers as these functions are candidates for a general role of sphingolipids within the environment, as opposed to taxon-specific associations.

More functions were enriched than depleted in potential sphingolipids producers throughout the water column, with few depleted functions overall (Fig. 5, Supplementary Fig. 6, Supplementary Data 4). This lack of negative associations suggests that the biosynthesis of sphingolipids does not consistently replace other functions in the Black Sea, and reflects the diverse roles of sphingolipids within bacteria. Enriched functions that were present in at least 75% of the sphingolipid producing phyla differed with depth (Fig. 5b,c), with transporter associations prevalent in the top part of the water column and associations with small molecule and nitrogen compound metabolic processes and transferases in the euxinic zone (Fig. 5c). Many enriched functions were related to enzymatic activity (Fig. 5c), suggesting a link between sphingolipids and microbial metabolism. Certain gene families with unknown functions were enriched in potential sphingolipid producers (Supplementary Data 4), which shows a lack of understanding of sphingolipid biosynthesis. Specific enriched functions were related to a response to oxidative stress and cell wall remodeling (Supplementary Data 4), possibly reflecting an adaptive role of sphingolipids in the euxinic zone of the Black Sea. These results provide a first and intriguing look into the role of sphingolipids in the Black Sea specifically and potentially in other environmental settings, pending future confirmation in laboratory cultures.

**Fig. 5.**
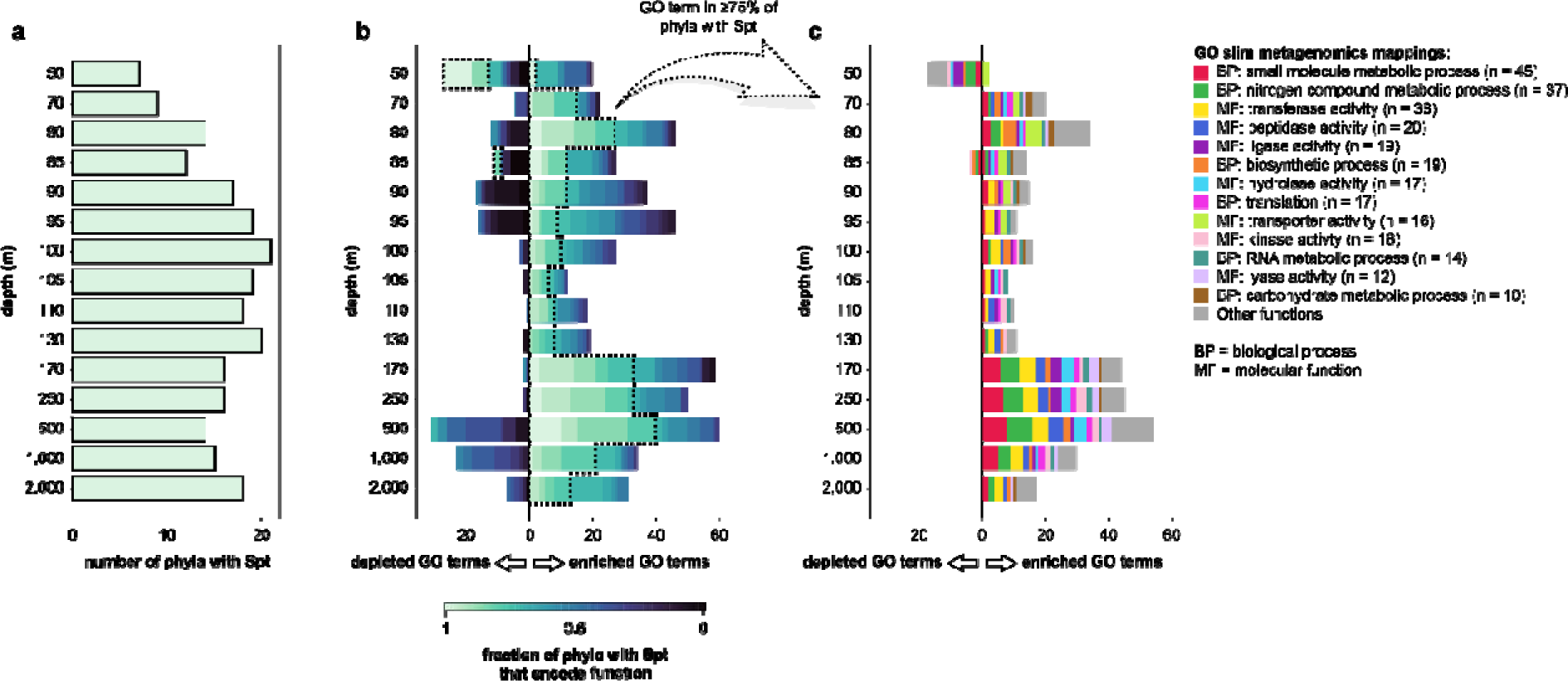
Pangenome-wide association analysis identifies functions that are depleted and enriched in MAGs that contain Spt-like proteins. (**a**) Number of phyla represented by the 704 MAGs that contain Spt-like proteins and whose (completeness – 5 x contamination) ≥ 70%. (**b**) Number of Gene Ontology (GO) terms that are depleted (to the left of 0) or enriched (to the right of 0) in MAGs containing Spt-like proteins compared to MAGs that do not contain Spt-like proteins based on the (qualitative) Fisher’s exact test. The test compares all MAGs that were binned from a specific depth in the water column. GO terms are colored according to the fraction of phyla with Spt-like proteins at that depth that contain at least one member that encodes the function. For the same figure for 5 other functional universes as well identification of depleted and enriched functions based on the (quantitative) Mann-Whitney U test see Supplementary Fig. 6. (**c**) Mapping of GO terms in panel a that are present in at least 75% of phyla that encode Spt-like proteins to GO slim metagenomics. GO terms can have multiple GO slim mappings. All depleted and enriched GO terms and their GO slim metagenomics mappings can be found in Supplementary Data 4.

## CONCLUSIONS

Recent advances in meta-omics technologies, including metagenomics, metatranscriptomics, metaproteomics, metabolomics, and metalipidomics, have revolutionized our capacity to characterize microbial communities in various environments^19, 73, 74^. However, to date, there have been few studies that have effectively combined metagenomics with metabolomics to elucidate the microbial taxa and their metabolites in complex settings^18, 19^. In an analysis of a global microbial community samples collected for the Earth Microbiome Project, Shaffer and colleagues^19^ demonstrated that metabolite diversity exhibits turnover and nestedness, related to both microbial communities and the environment. In our study, we present for the first time the combined application of integrated metalipidomics and metagenomics. This approach has been pivotal in studying a diverse array of lipids, leading to the identification of previously unknown sphingolipids and the prediction of their potential producers and functions in complex environmental matrices, and showing the potential of using this methodological approach in other environments. Further studies are needed to understand whether the novel sphingolipids detected here are taxonomically or environmentally specific, and their role in the cell in environmental settings.

## ONLINE METHODS

### Sampling

A detailed description of sample collection, DNA and lipid extraction, and UHPLC-HRMS/MS initial data processing is given in previous studies^7, 8, 28^. Briefly, suspended particulate matter (SPM) was collected in the water column of the Black Sea at 15 depths (50 to 2,000 mbsl) during the PHOXY cruise (9-10 June 2013). The PHOX2 sampling station was located at 42°53.8’N, 30°40.7’E in the western gyre of the Black Sea. SPM was collected with McLane WTS-LV in situ pumps (McLane Laboratories Inc., Falmouth) on pre-ashed 142 mm diameter 0.7 mm pore size glass fiber GF/F filters (Pall Corporation, Port Washington, NY). The filters were immediately stored at -80°C after collection.

### Lipid extraction

A modified Bligh-Dyer procedure was used to extract lipids from the freeze-dried filters. Lipids on the filters were extracted ultrasonically for 10 min, twice in a mixture of methanol, dichloromethane and phosphate buffer (2:1:0.8, v:v:v) and twice with a mixture of methanol, dichloromethane and aqueous trichloroacetic acid solution pH 3 (2:1:0.8, v:v:v). The organic phase was separated by adding additional dichloromethane and buffer to a final solvent ratio of 1:1:0.9 (v:v) and was re-extracted three times with dichloromethane and dried under a stream of N_2_ gas. The extract was redissolved in a mixture of MeOH:DCM (9:1, v:v) and filtered through 4 mm diameter 0.45 µm pore size regenerated cellulose syringe filters (Grace Alltech).

### UHPLC-HRMS/MS analysis and molecular networking

The lipid extracts were analyzed using an Agilent 1290 Infinity I Ultra High-Pressure Liquid Chromatography (UHPLC) system coupled to a Q Exactive Orbitrap high-resolution mass spectrometry (HRMS) system (Thermo Fisher Scientific, Waltham, MA). Instrument methods are described in a previous study^8^. The raw data files generated by UHPLC-HRMS/MS were further processed using MZmine software^75^. Processing steps^76^ included mass peak detection, chromatogram building, chromatogram deconvolution, isotope grouping, ion component alignment, gap filling, as well as a manual check.

The processed dataset of MS/MS spectra was further analyzed using the Feature Based Molecular Networking method^77^ through the Global Natural Product Social Molecular Networking (GNPS) platform^76^ to build molecular networks of the detected components. Molecular networking is a data analysis methodology utilized in untargeted metabolomics studies based on MS/MS analysis. It organizes MS/MS spectra into a network-like map, where molecules with similar spectral patterns are grouped together, indicating their structural similarity. Vector similarities were calculated by comparing pairs of spectra based on at least six matching fragment ions (peaks). This comparison takes into account the relative intensities of the fragment ions as well as the difference in precursor m/z values between the spectra^78^. The resulting molecular network is generated using MATLAB scripts, where each spectrum is allowed to connect to its top K scoring matches (typically up to 10 connections per node). Edges between spectra are retained if they are among the top K matches for both spectra and if the vector similarity score, represented as a cosine value, exceeds a user-defined threshold. In this study, a cosine value of 0.6 was used, with a cosine value of 1.0 indicating identical spectra. The exported molecular network was then imported into the visualization program Cytoscape^79, 80^ for further analysis.

Each node in the molecular network stands for an individual molecule associated with a specific MS/MS spectrum. Due to the extraction and analytical methods, and based on annotations of the same data in a previous study^7^, most of the ion components from the molecular network we detected were lipids, thus we used the term metalipidome where the ion components are discussed. Most of the molecules that clustered together in the subnetworks were either analogs of each other with an identical headgroup or with a similar core, differing by simple transformations such as alkylation, unsaturation, and glycosylation.

### Metalipidome annotation

The MS/MS spectra in the metalipidomic molecular network only resulted in <2% annotations based on a search through the GNPS library, as described in our previous work^7^. These 239 annotations included ca. 20 contaminants, and mostly TAGs, highly unsaturated PC-DAGs, PE-DAGs from plankton and algal species, leaving the vast majority of ion components unknown. Our previous work^7^ further annotated the most important lipids in the molecular network (with obvious changes of abundance through the water column or as a central node linked by several other lipids in the network) either by comparison to data from previous studies^10, 12–17, 28, 81–91^ or putatively identified based on accurate mass and MS/MS fragmentation. We submitted the mass spectra of these annotated lipids to the GNPS library for this study’s further analysis and for public use.

In total, the mass spectra of 249 representative lipids of the 18 major lipid classes from the molecular network were submitted to the GNPS library. Based on the annotated lipids from each subnetwork, we determined major lipid classes and used them for the lipid composition analysis.

### Lipid standard calibration

Calibration of lipid peak intensity using external standards is necessary to correct for large differences in ionization response between lipid classes, especially for the lipids with different polar headgroups. Supplementary Tables 1 and 2 detail the information on standards used to quantify each lipid class. We assumed that all lipids from the same class should have similar response factors, thus curves based on a single representative lipid species were used to calibrate each lipid class^92^. For sphingolipids Cers and LysCers, an average response factor of ceramide mixers was used for calibration (Supplementary Tables 1 and 2). 1-deoxyCer-associated sphingolipid classes were calibrated using an external standard 1-deoxyCer (d18:1/24:0) and Gly-Cers were calibrated using Glucosyl (β) C12 Cer. For the class of lipids with no standards available (e.g., OLs), a response factor from another class of amino lipids (DGTS) was used for calculation. The concentration of all the lipids of each MS/MS spectra (ng L^-1^) was then added as metadata for the metalipidome visualization across the water column of the Black Sea sample set (50– 2000 mbsl).

### DNA extraction

The DNA extraction from SPMs across the water column of the Black Sea was reported in a previous study^28, 93^. In short, DNA was extracted with the DNA Power Soil isolation kit (Mo Bio Laboratories, Inc., Carlsbad, CA). The integrity and concentration of the extracted DNA were tested by agarose gel electrophoresis and NanoDrop (Thermo Scientific, Waltham, MA) quantification.

### Metagenomic sequencing

The metagenomes of the samples used in this study were sequenced before in ref. ^93^. We sequenced the metagenomes again here, much deeper this time. TruSeq nano libraries were sequenced with Illumina MiSeq at Utrecht Sequencing Facility generating 251 base pair paired-end reads. All bioinformatic tools used to analyze these metagenomes were run with default parameters unless otherwise stated. FastQC v0.11.9 (https://www.bioinformatics.babraham.ac.uk/projects/fastqc/) was used for quality control, and quality and adapter trimming was performed with Trimmomatic v0.39 (ref. ^94^) with options “ILLUMINACLIP:TruSeq3-PE-2.fa:2:30:12:2:TRUE MAXINFO:40:0.6 MINLEN:40”. After trimming, the samples contained 197,509,556 - 412,252,887 paired-end sequencing reads.

### Assembly and binning

The 15 samples were individually assembled with metaSPAdes v3.15.1(ref. ^95^) and assembly quality was assessed with QUAST v5.0.2 (ref. ^96^). Sequencing reads were mapped back to the assembled scaffolds with BWA-MEM v2.2.1 (ref. ^97^) generating 15 x 15 mappings, and the scaffolds were binned per sample with MetaBAT2 v2.2.15 (ref. ^98^), CONCOCT v1.1.0 (ref. ^99^), and MaxBin2 v2.2.7 (ref. ^100^). For CONCOCT, the scaffolds were cut in chunks of 10,000 base pairs, and scaffolds shorter than 2,500 base pairs were excluded from binning. For MaxBin2, scaffolds shorter than 2,500 base pairs were excluded from binning and the 40 universal marker genes set was used. Metagenome-assembled genomes (MAGs) generated by MetaBAT2, CONCOCT, and MaxBin2 were used as input for DAS Tool v1.1.3 (ref. ^101^) with the BLAST search engine to arrive at a final MAG set per sample. MAG quality was assessed with CheckM v1.1.3 (ref. ^102^) in the lineage-specific workflow.

Taxonomy was assigned to scaffolds and MAGs with medium to high quality (>50% completeness, <10% contamination^65^) with Contig Annotation Tool (CAT) and Bin Annotation Tool (BAT)^103^, respectively, from the CAT pack software suite v5.2.3. A reference database was constructed based on the set of non-redundant proteins in GTDB r207 (ref. ^104^). Proteins were predicted with Prodigal v2.6.3 in metagenomic mode^105^ and aligned to the reference database with DIAMOND v2.0.6 (ref. ^106^) with the –top parameter set to 11 for CAT and the –top parameter set to 6 for BAT. With BAT its default parameters (-f 0.3), a MAG may get multiple taxonomic annotations. In those cases, the majority classification (-f 0.5) was used.

### Homology searches of ceramide biosynthesis genes and phylogeny of Spt homologs

Genes were predicted on the scaffolds with Prodigal v2.6.3 in metagenomic mode^105^. We queried the bacterial serine palmitoyltransferase (Spt), bacterial ceramide synthase (bCerS), and ceramide reductase (CerR) protein sequences from ref. ^37^ (query sequences and TBLASTN results from Supplementary Data 1 of ref. ^37^ in the predicted protein sequences with BLASTP v2.12.0 (ref. ^37^), with a database size of 1 x 10^8^ to make *e*-values comparable across samples. Spt homologs with an *e*-value ≤ 1 x 10^-70^ and query coverage ≥ 70% to the best hit query sequence based on *e*-value were selected for further phylogenetic tree building with known alpha-oxoamine synthase sequences to constrain Spt activity. Some of these hits were part of longer protein sequences. To prevent alignment of possible non-homologous regions, we extracted the BLAST High Scoring Pair (HSP) from hits with a sequence coverage < 70%, and in other cases the entire predicted protein. The hits were clustered together with the Spt query sequences and 16 alpha-oxoamine synthases with different enzymatic activities^6^ with CD-HIT v4.8.1 (ref. ^107^) at 90% sequence identity. The resulting 2,666 representative sequences were aligned with MAFFT v7.505 using the L-INS-i algorithm, and the alignment was trimmed with trimAl v1.4.rev15 (ref. ^108^) in gappyout mode, leaving 411 sites. A maximum likelihood phylogenetic tree was reconstructed with IQ-TREE v2.1.2 (ref. ^109^) using model finder^110^ with nuclear amino-acid models, and 1,000 ultrafast bootstraps^111^. The LG+F+R10 model was chosen based on Bayesian Information Criterion (BIC). The tree was visualized in iTOL^112^. Homologs of bCerS and CerR were filtered based on *e*-value ≤ 1 x 10^-30^ and query coverage ≥ 50%.

To identify possible eukaryotic Spt sequences, we queried the Spt sequences of *Homo sapiens* (NP_006406.1), *Arabidopsis thaliana* (NP_190447.1), and *Saccharomyces cerevisiae* (CAA56805.1) in the predicted protein sequences with BLASTP, with a database size of 1 x 10^8^, and filtered based on *e*-value ≤ 1 x 10^-70^ and query coverage ≥ 70%.

### Abundance estimation

Taxonomic profiles were generated based on mapping of the sequencing reads to the scaffolds with BWA-MEM v2.2.1 (ref. ^97^). Taxonomic annotation of a read was inherited from the BAT annotation of the MAG when possible or else from the CAT annotation of the assembled scaffold, similar to ref. ^66^. We similarly generated profiles for the mappings that could be associated with Spt-like proteins via the MAG or unbinned scaffold.

### Pangenome-wide association analyses

The MAGs whose (completeness – 5 x contamination) ≥ 70% were annotated in 6 functional universes provided by InterPro^113, 114^ (Conserved Domains Database (CDD), Gene Ontology (GO), InterProScan (IPR), NCBIfam, pathways, and Pfam) with InterProScan v5.63-95.0 (ref. ^113, 114^). In each SPM sample, functions that were more commonly present or absent (qualitatively) in potential sphingolipid producers compared to MAGs that lack Spt-like proteins were identified with Fisher’s exact test, using Bonferroni *p*-value correction. Functions that were present in higher or lower abundance (quantitatively) were identified with the Mann-Whitney U test, using Bonferroni *p*-value correction. We performed the tests per SPM sample to allow for the identification of niche-specific associations.

### Network analyses and statistics

Spearman’s correlation coefficient between the abundance of scaffolds that contain *Spt* genes and the abundance of sphingolipids across the water column of the Black Sea were calculated with Python (version 3.97). Significant positive correlations (r ≥ 0.9, *P* ≤ 0.001) were selected to build the sphingolipid and their potential producers’ network^115^ using Cytoscape (version 3.9.0) with a Spring Embedded layout^79, 80^. Scaffolds and sphingolipids were set as nodes in the network, and edges with a Spearman correlation coefficient > 0.9 were shown.

In order to reduce this data sparsity, thresholds of 1 x 10^6^ for sphingolipid peak intensity and 1 for scaffold read among 15 samples were used to filter the datasets. A total of 1,455 scaffolds and 229 sphingolipid species were selected to build the positive network. 133 sphingolipid structures had a Spearman rank correlation coefficient ≥ 0.9 with 728 binned scaffolds and were visualized in a network.

To represent the abundance weight of scaffolds and sphingolipids across the water column of the Black Sea (110-2,000 mbsl), a color gradient stripe ranging from orange to navy blue was employed. The abundance weight of the ith MS/MS and scaffolds among the samples was calculated using the following formula:

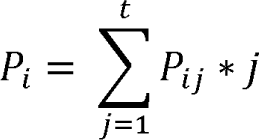

where *P_ij_* corresponds to the relative frequency of the ith MS/MS or scaffolds (i = 1, 2, …, m) in the jth sample (j = 1, 2, …, t), to illustrate the dominant distribution of a specific sphingolipid or scaffold across the water column. For instance, when *P_ij_* has a value close to 9, it represents a sphingolipid or scaffold that exhibits significant abundance in the upper euxinic zone (∼110 mbsl). Such instances are denoted by the color orange on the gradient stripe. Conversely, when *P_ij_* approaches a value of 15, it indicates a predominant distribution in the deep waters (∼2000 mbsl), and is consequently represented by the color dark blue on the stripe. By examining the network, a trend can be observed from the upper euxinic zone to the deep sea in terms of abundance levels.

## ACKNOWLEDGEMENT

We acknowledge Denise Dorhout and Monique Verweij for the lipid standard calibration. We thank Diana X. Sahonero-Canavesi and Toby Halamka for the discussion of sphingolipids and their genes. We thank Saara Suominen, Zhao Zhao, Nina Dombrowski, Pierre Ramond, Julia C. Engelmann, and Milou Arts for their discussions on the omics network. We thank Pierre Offre for suggestions on the pan-genome-wide association analyses. We also thank Alejandro Abdala Asbun, Sanne Vreugdenhil, Maartje Brouwer, Anchelique Mets and Jort Ossebaar for their technical support. Ellen C. Hopmans and Stefan Schouten provided valuable comments for the initial draft. LV and JSSD received funding from the Soehngen Institute for Anaerobic Microbiology (SIAM) through a Gravitation Grant (024.002.002) from the Dutch Ministry of Education, Culture, and Science (OCW). JSSD received funding from the European Research Council (ERC) under the European Union’s Horizon 2020 research and innovation program (grant agreement no. 694569—MICROLIPIDS) and from a Spinoza award from NWO.

The authors declare no conflict of interest.

## AUTHORS CONTRIBUTIONS

S.D., F.A.B.v.M., N.J.B., J.S.S.D., and L.V. conceived the study. S.D. performed metalipidome data analyses. F.A.B.v.M. performed metagenome and pangenome data analyses. S.D. and N.J.B. performed lipid identifications. S.D. and F.A.B.v.M. performed sphingolipids-microbiome association analyses. S.D. and F.A.B.v.M. wrote the draft manuscript. J.S.S.D. and L.V. supervised the study. All authors read, discussed, and approved the final version of the manuscript.

## DATA AVAILABILITY

The processed data (.mgf and .csv) with the molecular network and detailed parameter settings can be accessed at the GNPS platform: https://gnps.ucsd.edu/ProteoSAFe/status.jsp?task=2e274c97c13e4f7d8e38637ffa9f0bee.

249 representative lipids of the 18 major lipid classes from the water column of the Black Sea were submitted to the GNPS library:

https://gnps.ucsd.edu/ProteoSAFe/status.jsp?task=dc888b40c9c848fc9ece1738e9e434f0.

The metagenomic reads are uploaded at XXXX under accession XXXX, the assemblies at XXXX under accession XXXX, and the MAGs at XXXX under accession XXXX.

The source data of figures are available at Zenodo at DOI: 10.5281/zenodo.10346514.

## CODE AVAILABILITY

All codes used in this study are available at Zenodo at DOI: 10.5281/zenodo.10346514.

## REFERENCES

1. Sohlenkamp, C. & Geiger, O. Bacterial membrane lipids: diversity in structures and pathways. FEMS Microbiology Reviews 40, 133–159 (2016).

2. Ding, S. et al. Changes in the membrane lipid composition of a Sulfurimonas species depend on the electron acceptor used for sulfur oxidation. ISME Communications 2, 121 (2022).

3. Van Mooy, B.A.S. et al. Phytoplankton in the ocean use non-phosphorus lipids in response to phosphorus scarcity. Nature 458, 69–72 (2009).

4. Weijers, J.W.H. et al. Membrane lipids of mesophilic anaerobic bacteria thriving in peats have typical archaeal traits. Environmental Microbiology 8, 648–657 (2006).

5. Geiger, O., López-Lara, I.M. & Sohlenkamp, C. Phosphatidylcholine biosynthesis and function in bacteria. Biochimica et Biophysica Acta (BBA) - Molecular and Cell Biology of Lipids 1831, 503–513 (2013).

6. Geiger, O., González-Silva, N., López-Lara, I.M. & Sohlenkamp, C. Amino acid-containing membrane lipids in bacteria. Progress in Lipid Research 49, 46–60 (2010).

7. Ding, S., et al. Lipidomics of Environmental Microbial Communities. II: Characterization Using Molecular Networking and Information Theory. Frontiers in Microbiology 12 (2021).

8. Bale, N.J. et al. Lipidomics of Environmental Microbial Communities. I: Visualization of Component Distributions Using Untargeted Analysis of High-Resolution Mass Spectrometry Data. Frontiers in Microbiology 12 (2021).

9. Wakeham, S.G. et al. Biomarkers, chemistry and microbiology show chemoautotrophy in a multilayer chemocline in the Cariaco Basin. Deep Sea Research Part I: Oceanographic Research Papers 63, 133–156 (2012).

10. Popendorf, K.J. et al. Gradients in intact polar diacylglycerolipids across the Mediterranean Sea are related to phosphate availability. Biogeosciences 8, 3733–3745 (2011).

11. Van Mooy, B.A.S. & Fredricks, H.F. Bacterial and eukaryotic intact polar lipids in the eastern subtropical South Pacific: Water-column distribution, planktonic sources, and fatty acid composition. Geochimica et Cosmochimica Acta 74, 6499–6516 (2010).

12. Schubotz, F., Wakeham, S.G., Lipp, J.S., Fredricks, H.F. & Hinrichs, K.-U. Detection of microbial biomass by intact polar membrane lipid analysis in the water column and surface sediments of the Black Sea. Environmental Microbiology 11, 2720–2734 (2009).

13. Wakeham, S.G. et al. Microbial ecology of the stratified water column of the Black Sea as revealed by a comprehensive biomarker study. Organic Geochemistry 38, 2070–2097 (2007).

14. Schubotz, F., Xie, S., Lipp, J.S., Hinrichs, K.U. & Wakeham, S.G. Intact polar lipids in the water column of the eastern tropical North Pacific: abundance and structural variety of non-phosphorus lipids. Biogeosciences 15, 6481–6501 (2018).

15. Bale, N.J. et al. Impact of trophic state on the distribution of intact polar lipids in surface waters of lakes. Limnology and Oceanography 61, 1065–1077 (2016).

16. Ding, S. et al. Characteristics and origin of intact polar lipids in soil organic matter. Soil Biology and Biochemistry 151, 108045 (2020).

17. Liu, X.-L. et al. Identification of polar lipid precursors of the ubiquitous branched GDGT orphan lipids in a peat bog in Northern Germany. Organic Geochemistry 41, 653–660 (2010).

18. Thukral, M., Allen, A.E. & Petras, D. Progress and challenges in exploring aquatic microbial communities using non-targeted metabolomics. The ISME Journal 17, 2147–2159 (2023).

19. Shaffer, J.P. et al. Standardized multi-omics of Earth’s microbiomes reveals microbial and metabolite diversity. Nature Microbiology 7, 2128–2150 (2022).

20. Wakeham, S.G. & Lee, C. Limits of our knowledge, part 2: Selected frontiers in marine organic biogeochemistry. Marine Chemistry 212, 16–46 (2019).

21. Steen, A.D., et al. Analytical and Computational Advances, Opportunities, and Challenges in Marine Organic Biogeochemistry in an Era of “Omics”. Frontiers in Marine Science 7 (2020).

22. Pearson, A., Flood Page, S.R., Jorgenson, T.L., Fischer, W.W. & Higgins, M.B. Novel hopanoid cyclases from the environment. Environmental Microbiology 9, 2175–2188 (2007).

23. Pearson, A. et al. Diversity of hopanoids and squalene-hopene cyclases across a tropical land-sea gradient. Environmental Microbiology 11, 1208–1223 (2009).

24. Pérez Gallego, R., Bale, N.J., Sinninghe Damste, J.S. & Villanueva, L. Developing a genetic approach to target cyanobacterial producers of heterocyte glycolipids in the environment. Frontiers in Microbiology 14 (2023).

25. Kebschull, J.M. & Zador, A.M. Sources of PCR-induced distortions in high-throughput sequencing data sets. Nucleic Acids Research 43, e143–e143 (2015).

26. Jansson, J.K. & Baker, E.S. A multi-omic future for microbiome studies. Nature Microbiology 1, 16049 (2016).

27. Hauptfeld, E. et al. A metagenomic portrait of the microbial community responsible for two decades of bioremediation of poly-contaminated groundwater. Water Research 221, 118767 (2022).

28. Sollai, M., Villanueva, L., Hopmans, E.C., Reichart, G.-J. & Sinninghe Damsté, J.S. A combined lipidomic and 16S rRNA gene amplicon sequencing approach reveals archaeal sources of intact polar lipids in the stratified Black Sea water column. Geobiology 17, 91–109 (2019).

29. Dijkstra, N. et al. Phosphorus dynamics in and below the redoxcline in the Black Sea and implications for phosphorus burial. Geochimica et Cosmochimica Acta 222, 685–703 (2018).

30. Hu, Q. et al. Microalgal triacylglycerols as feedstocks for biofuel production: perspectives and advances. The Plant Journal 54, 621–639 (2008).

31. Geiger, O., Padilla-Gómez, J. & López-Lara, I.M. Bacterial Sphingolipids and Sulfonolipids, in Biogenesis of Fatty Acids, Lipids and Membranes. (ed. O. Geiger) 123–137 (Springer International Publishing, Cham; 2019).

32. Harrison, P.J., Dunn, T. & Campopiano, D.J. Sphingolipid biosynthesis in man and microbes. Natural Product Reports 35, 921–954 (2018).

33. Nelson, D.L. & Cox, M.M. Lehninger Principles of Biochemistry. 7th Edition. (W.H. Freeman, New York; 2017).

34. Schleyer, G. et al. Lipid biomarkers for algal resistance to viral infection in the ocean. Proceedings of the National Academy of Sciences 120, e2217121120 (2023).

35. Schleyer, G. et al. In plaque-mass spectrometry imaging of a bloom-forming alga during viral infection reveals a metabolic shift towards odd-chain fatty acid lipids. Nature Microbiology 4, 527–538 (2019).

36. Ziv, C. et al. Viral serine palmitoyltransferase induces metabolic switch in sphingolipid biosynthesis and is required for infection of a marine alga. Proceedings of the National Academy of Sciences 113, E1907–E1916 (2016).

37. Stankeviciute, G. et al. Convergent evolution of bacterial ceramide synthesis. Nature Chemical Biology 18, 305–312 (2022).

38. Muñoz-Garcia, A., Ro, J., Brown, J.C. & Williams, J.B. Identification of complex mixtures of sphingolipids in the stratum corneum by reversed-phase high-performance liquid chromatography and atmospheric pressure photospray ionization mass spectrometry. Journal of Chromatography A 1133, 58–68 (2006).

39. Hannich, J.T. et al. 1-Deoxydihydroceramide causes anoxic death by impairing chaperonin-mediated protein folding. Nature Metabolism 1, 996–1008 (2019).

40. Duan, J. & Merrill, A.H., Jr. 1-Deoxysphingolipids encountered exogenously and made de novo: Dangerous mysteries inside an enigma. The Journal of biological chemistry 290, 15380–15389 (2015).

41. Cuadros, R. et al. The marine compound spisulosine, an inhibitor of cell proliferation, promotes the disassembly of actin stress fibers. Cancer Letters 152, 23–29 (2000).

42. Zitomer, N.C. et al. Ceramide synthase inhibition by fumonisin B1 causes accumulation of 1-deoxysphinganine: a novel category of bioactive 1-deoxysphingoid bases and 1-deoxydihydroceramides biosynthesized by mammalian cell lines and animals. Journal of Biological Chemistry 284, 4786–4795 (2009).

43. Pruett, S.T. et al. Thematic Review Series: Sphingolipids. Biodiversity of sphingoid bases (“sphingosines”) and related amino alcohols. Journal of Lipid Research 49, 1621–1639 (2008).

44. Sandhoff, R. Very long chain sphingolipids: Tissue expression, function and synthesis. FEBS Letters 584, 1907–1913 (2010).

45. Furland, N.E., Zanetti, S.R., Oresti, G.M., Maldonado, E.N. & Aveldaño, M.I. Ceramides and sphingomyelins with high proportions of very long-chain polyunsaturated fatty acids in mammalian germ cells. Journal of Biological Chemistry 282, 18141–18150 (2007).

46. Hannun, Y.A. & Obeid, L.M. Sphingolipids and their metabolism in physiology and disease. Nature Reviews Molecular Cell Biology 19, 175–191 (2018).

47. Alecu, I. et al. Localization of 1-deoxysphingolipids to mitochondria induces mitochondrial dysfunction. Journal of Lipid Research 58, 42–59 (2017).

48. Zanetti, S.R., de los Ángeles Monclus, M., Rensetti, D.E., Fornés, M.W. & Aveldaño, M.I. Ceramides with 2-hydroxylated, very long-chain polyenoic fatty acids in rodents: From testis to fertilization-competent spermatozoa. Biochimie 92, 1778–1786 (2010).

49. Vasireddy, V. et al. Loss of functional ELOVL4 depletes very long-chain fatty acids (≥C28) and the unique ω-O-acylceramides in skin leading to neonatal death. Human Molecular Genetics 16, 471–482 (2007).

50. Michaelson, L.V., Napier, J.A., Molino, D. & Faure, J.-D. Plant sphingolipids: Their importance in cellular organization and adaption. Biochimica et Biophysica Acta (BBA) - Molecular and Cell Biology of Lipids 1861, 1329–1335 (2016).

51. Luttgeharm, K.D., Kimberlin, A.N. & Cahoon, E.B. Plant sphingolipid metabolism and function, in Lipids in Plant and Algae Development. (eds. Y. Nakamura & Y. Li-Beisson) 249–286 (Springer International Publishing, Cham; 2016).

52. Markham, J.E. et al. Sphingolipids containing very-long-chain fatty acids define a secretory pathway for specific polar plasma membrane protein targeting in Arabidopsis The Plant Cell 23, 2362–2378 (2011).

53. Ternes, P. et al. Disruption of the ceramide synthase LOH1 causes spontaneous cell death in Arabidopsis thaliana. New Phytologist 192, 841–854 (2011).

54. Roudier, F. et al. Very-long-chain fatty acids are involved in polar auxin transport and developmental patterning in Arabidopsis The Plant Cell 22, 364–375 (2010).

55. Menuz, V. et al. Protection of C. elegans from anoxia by HYL-2 ceramide synthase. Science 324, 381–384 (2009).

56. Schneiter, R. et al. Electrospray ionization tandem mass spectrometry (ESI-MS/MS) analysis of the lipid molecular species composition of yeast subcellular membranes reveals acyl chain-based sorting/remodeling of distinct molecular species en route to the plasma membrane. Journal of Cell Biology 146, 741–754 (1999).

57. Aguilera-Romero, A., Gehin, C. & Riezman, H. Sphingolipid homeostasis in the web of metabolic routes. Biochimica et Biophysica Acta (BBA) - Molecular and Cell Biology of Lipids 1841, 647–656 (2014).

58. Guillas, I. et al. C26-CoA-dependent ceramide synthesis of Saccharomyces cerevisiae is operated by Lag1p and Lac1p. EMBO Journal 20, 2655–2665 (2001).

59. Vardi, A. et al. Viral glycosphingolipids induce lytic infection and cell death in marine phytoplankton. Science 326, 861–865 (2009).

60. Řezanka, T. & Sigler, K. Odd-numbered very-long-chain fatty acids from the microbial, animal and plant kingdoms. Progress in Lipid Research 48, 206–238 (2009).

61. Ikushiro, H., Hayashi, H. & Kagamiyama, H. A water-soluble homodimeric serine palmitoyltransferase from Sphingomonas paucimobilis EY2395T strain: Purification, characterization, cloning, and overproduction. Journal of Biological Chemistry 276, 18249–18256 (2001).

62. Yard, B.A. et al. The structure of serine palmitoyltransferase; gateway to sphingolipid biosynthesis. Journal of Molecular Biology 370, 870–886 (2007).

63. Brown, E.M. et al. Bacteroides-derived sphingolipids are critical for maintaining intestinal homeostasis and symbiosis. Cell Host & Microbe 25, 668–680.e667 (2019).

64. Olea-Ozuna, R.J. et al. Five structural genes required for ceramide synthesis in Caulobacter and for bacterial survival. Environmental Microbiology 23, 143–159 (2021).

65. Bowers, R.M. et al. Minimum information about a single amplified genome (MISAG) and a metagenome-assembled genome (MIMAG) of bacteria and archaea. Nature Biotechnology 35, 725–731 (2017).

66. Ernestina, H. et al. Integration of taxonomic signals from MAGs and contigs improves read annotation and taxonomic profiling of metagenomes. bioRxiv, 2023.2003.2022.533753 (2023).

67. Bertagnolli, A.D., Padilla, C.C., Glass, J.B., Thamdrup, B. & Stewart, F.J. Metabolic potential and in situ activity of marine Marinimicrobia bacteria in an anoxic water column. Environmental Microbiology 19, 4392–4416 (2017).

68. Hawley, A.K. et al. Diverse Marinimicrobia bacteria may mediate coupled biogeochemical cycles along eco-thermodynamic gradients. Nature Communications 8, 1507 (2017).

69. Lewis, W.H., Tahon, G., Geesink, P., Sousa, D.Z. & Ettema, T.J.G. Innovations to culturing the uncultured microbial majority. Nature Reviews Microbiology 19, 225–240 (2021).

70. Liang, B. et al. Anaerolineaceae and Methanosaeta turned to be the dominant microorganisms in alkanes-dependent methanogenic culture after long-term of incubation. AMB Express 5, 37 (2015).

71. Parkes, R.J., Gibson, G.R., Mueller-Harvey, I., Buckingham, W.J. & Herbert, R.A. Determination of the substrates for sulphate-reducing bacteria within marine and esturaine sediments with different rates of sulphate reduction. Microbiology 135, 175–187 (1989).

72. Fonseca, A., Espinoza, C., Nielsen, L.P., Marshall, I.P.G. & Gallardo, V.A. Bacterial community of sediments under the Eastern Boundary Current System shows high microdiversity and a latitudinal spatial pattern. Frontiers in Microbiology 13 (2022).

73. Biswapriya, B.M., Carl, L., Michael, O. & Laura, A.C. Integrated omics: tools, advances and future approaches. Journal of Molecular Endocrinology 62, R21–R45 (2019).

74. Revel, M., Châtel, A. & Mouneyrac, C. Omics tools: New challenges in aquatic nanotoxicology? Aquatic Toxicology 193, 72–85 (2017).

75. Pluskal, T., Castillo, S., Villar-Briones, A. & Orešič, M. MZmine 2: Modular framework for processing, visualizing, and analyzing mass spectrometry-based molecular profile data. BMC Bioinformatics 11, 395 (2010).

76. Wang, M. et al. Sharing and community curation of mass spectrometry data with Global Natural Products Social Molecular Networking. Nature Biotechnology 34, 828–837 (2016).

77. Nothias, L.-F. et al. Feature-based molecular networking in the GNPS analysis environment. Nature Methods 17, 905–908 (2020).

78. Watrous, J. et al. Mass spectral molecular networking of living microbial colonies. Proceedings of the National Academy of Sciences 109, E1743 (2012).

79. Smoot, M.E., Ono, K., Ruscheinski, J., Wang, P.-L. & Ideker, T. Cytoscape 2.8: new features for data integration and network visualization. Bioinformatics 27, 431–432 (2010).

80. Shannon, P. et al. Cytoscape: A Software Environment for Integrated Models of Biomolecular Interaction Networks. Genome Research 13, 2498–2504 (2003).

81. Becker, K.W. et al. Combined pigment and metatranscriptomic analysis reveals highly synchronized diel patterns of phenotypic light response across domains in the open oligotrophic ocean. The ISME Journal (2020).

82. Becker, K.W. et al. Isoprenoid quinones resolve the stratification of redox processes in a biogeochemical continuum from the photic zone to deep anoxic sediments of the Black Sea. Applied and Environmental Microbiology 84, e02736–02717 (2018).

83. Kharbush, J.J., Allen, A.E., Moustafa, A., Dorrestein, P.C. & Aluwihare, L.I. Intact polar diacylglycerol biomarker lipids isolated from suspended particulate organic matter accumulating in an ultraoligotrophic water column. Organic Geochemistry 100, 29–41 (2016).

84. Wörmer, L., Lipp, J.S. & Hinrichs, K.-U. Comprehensive analysis of microbial lipids in environmental samples through HPLC-MS protocols, in Hydrocarbon and Lipid Microbiology Protocols: Petroleum, Hydrocarbon and Lipid Analysis. (eds. T.J. McGenity, K.N. Timmis & B. Nogales) 289–317 (Springer Berlin Heidelberg, Berlin, Heidelberg; 2015).

85. Moore, E.K. et al. Novel mono-, di-, and trimethylornithine membrane lipids in northern wetland Planctomycetes. Applied and Environmental Microbiology 79, 6874 (2013).

86. Ding, S. et al. In situ production of core and intact bacterial and archaeal tetraether lipids in groundwater. Organic Geochemistry 126, 1–12 (2018).

87. Wörmer, L. et al. A micrometer-scale snapshot on phototroph spatial distributions: mass spectrometry imaging of microbial mats in Octopus Spring, Yellowstone National Park. Geobiology 18, 742–759 (2020).

88. Liu, X.-L., Summons, R.E. & Hinrichs, K.-U. Extending the known range of glycerol ether lipids in the environment: structural assignments based on tandem mass spectral fragmentation patterns. Rapid Communications in Mass Spectrometry 26, 2295–2302 (2012).

89. Cui, X. et al. Niche expansion for phototrophic sulfur bacteria at the Proterozoic–Phanerozoic transition. Proceedings of the National Academy of Sciences, 202006379 (2020).

90. Welti, R. & Wang, X. Lipid species profiling: a high-throughput approach to identify lipid compositional changes and determine the function of genes involved in lipid metabolism and signaling. Current Opinion in Plant Biology 7, 337–344 (2004).

91. Sturt, H.F., Summons, R.E., Smith, K., Elvert, M. & Hinrichs, K.-U. Intact polar membrane lipids in prokaryotes and sediments deciphered by high-performance liquid chromatography/electrospray ionization multistage mass spectrometry—new biomarkers for biogeochemistry and microbial ecology. Rapid Communications in Mass Spectrometry 18, 617–628 (2004).

92. Holm Henry, C., et al. Global ocean lipidomes show a universal relationship between temperature and lipid unsaturation. Science 376, 1487–1491 (2022).

93. Villanueva, L. et al. Bridging the membrane lipid divide: bacteria of the FCB group superphylum have the potential to synthesize archaeal ether lipids. The ISME Journal 15, 168–182 (2021).

94. Bolger, A.M., Lohse, M. & Usadel, B. Trimmomatic: a flexible trimmer for Illumina sequence data. Bioinformatics 30, 2114–2120 (2014).

95. Nurk, S., Meleshko, D., Korobeynikov, A. & Pevzner, P.A. metaSPAdes: a new versatile metagenomic assembler. Genome Research 27, 824–834 (2017).

96. Gurevich, A., Saveliev, V., Vyahhi, N. & Tesler, G. QUAST: quality assessment tool for genome assemblies. Bioinformatics 29, 1072–1075 (2013).

97. Li, H. Aligning sequence reads, clone sequences and assembly contigs with BWA-MEM. arXiv preprint arXiv:1303.3997 (2013).

98. Kang, D.D., Froula, J., Egan, R. & Wang, Z. MetaBAT, an efficient tool for accurately reconstructing single genomes from complex microbial communities. PeerJ 3, e1165 (2015).

99. Alneberg, J. et al. Binning metagenomic contigs by coverage and composition. Nature Methods 11, 1144–1146 (2014).

100. Wu, Y.-W., Simmons, B.A. & Singer, S.W. MaxBin 2.0: an automated binning algorithm to recover genomes from multiple metagenomic datasets. Bioinformatics 32, 605–607 (2016).

101. Sieber, C.M.K. et al. Recovery of genomes from metagenomes via a dereplication, aggregation and scoring strategy. Nature Microbiology 3, 836–843 (2018).

102. Parks, D.H., Imelfort, M., Skennerton, C.T., Hugenholtz, P. & Tyson, G.W. CheckM: assessing the quality of microbial genomes recovered from isolates, single cells, and metagenomes. Genome Research 25, 1043–1055 (2015).

103. von Meijenfeldt, F.A.B., Arkhipova, K., Cambuy, D.D., Coutinho, F.H. & Dutilh, B.E. Robust taxonomic classification of uncharted microbial sequences and bins with CAT and BAT. Genome Biology 20, 217 (2019).

104. Parks, D.H. et al. GTDB: an ongoing census of bacterial and archaeal diversity through a phylogenetically consistent, rank normalized and complete genome-based taxonomy. Nucleic Acids Research 50, D785–D794 (2022).

105. Hyatt, D., LoCascio, P.F., Hauser, L.J. & Uberbacher, E.C. Gene and translation initiation site prediction in metagenomic sequences. Bioinformatics 28, 2223–2230 (2012).

106. Buchfink, B., Reuter, K. & Drost, H.-G. Sensitive protein alignments at tree-of-life scale using DIAMOND. Nature Methods 18, 366–368 (2021).

107. Fu, L., Niu, B., Zhu, Z., Wu, S. & Li, W. CD-HIT: accelerated for clustering the next-generation sequencing data. Bioinformatics 28, 3150–3152 (2012).

108. Capella-Gutiérrez, S., Silla-Martínez, J.M. & Gabaldón, T. trimAl: a tool for automated alignment trimming in large-scale phylogenetic analyses. Bioinformatics 25, 1972–1973 (2009).

109. Nguyen, L.-T., Schmidt, H.A., von Haeseler, A. & Minh, B.Q. IQ-TREE: A fast and effective stochastic algorithm for estimating maximum-likelihood phylogenies. Molecular Biology and Evolution 32, 268–274 (2015).

110. Kalyaanamoorthy, S., Minh, B.Q., Wong, T.K.F., von Haeseler, A. & Jermiin, L.S. ModelFinder: fast model selection for accurate phylogenetic estimates. Nature Methods 14, 587–589 (2017).

111. Hoang, D.T., Chernomor, O., von Haeseler, A., Minh, B.Q. & Vinh, L.S. UFBoot2: Improving the Ultrafast Bootstrap Approximation. Molecular Biology and Evolution 35, 518–522 (2018).

112. Letunic, I. & Bork, P. Interactive Tree Of Life (iTOL) v5: an online tool for phylogenetic tree display and annotation. Nucleic Acids Research 49, W293–W296 (2021).

113. Blum, M. et al. The InterPro protein families and domains database: 20 years on. Nucleic Acids Research 49, D344–D354 (2021).

114. Jones, P. et al. InterProScan 5: genome-scale protein function classification. Bioinformatics 30, 1236–1240 (2014).

115. Zhao, Z. et al. Microbial transformation of virus-induced dissolved organic matter from picocyanobacteria: coupling of bacterial diversity and DOM chemodiversity. The ISME Journal 13, 2551–2565 (2019).

