## Supplementary Information for "Coupled metalipidomics-metagenomics reveal structurally diverse sphingolipids produced by a wide variety of marine bacteria"

**Supplementary materials**

**SUPPLEMENTARY NOTE**

**Background on sphingolipids**

Sphingolipids are composed of a sphingosine backbone linked to a fatty acid via an amide bond. They play essential roles in eukaryotes as fundamental building blocks of the cell membrane and serve vital functions in cellular signaling and organization of lipid rafts^1-3^, where diverse sphingolipids are for example implicated in human disease^4-7^, plant pollen development^8, 9^, and response to environmental stresses^10, 11^. Despite their ubiquity and multifunctionality in eukaryotes, sphingolipids were thought to be rare in bacteria. The bacterial species that were first observed to produce them were given the prefix ‘Sphingo’ in their generic name to indicate this significance, i.e. *Sphingomonas* (phylum *Proteobacteria*) and *Sphingobacterium* (phylum *Bacteroidota*) ^1^. Over time, more sphingolipid-producing bacteria have been found, including many members of the phylum *Bacteroidota*, and a few members of the phyla *Chlorobi*, *Proteobacteria*, *Myxococcota*, and *Bdellovibrionota*^12, 13^. The metabolic pathway leading to the biosynthesis of the most common sphingolipid subtype and likely precursor of other sphingolipids, ceramide, has recently been elucidated in the bacterium *Caulobacter crescentus*^14^, and homology searches of key ceramide biosynthesis genes throughout the bacterial domain revealed potential sphingolipid production among 9 bacterial phyla (see Supplementary Data 2 for the taxonomic annotation of these sequences based on the Genome Taxonomy Database).

The functions of sphingolipids in the bacterial domain remain largely unknown and are likely diverse. Sphingolipids have been shown to replace the lipopolysaccharides of the bacterial outer membrane^15, 16^, but sphingolipid biosynthesis genes have also been identified in Gram-positive bacteria that lack this outer membrane^14^. Sphingolipids are known to accumulate during fruiting body formation in *Myxococcus xanthus*^17^. Parasitic *Bacteriovorax stolpii* that cannot synthesize sphingolipids exhibit a reduced ability to penetrate host cells^18^. Additionally, *Caulobacter crescentus*, when grown under phosphate-limiting conditions, increases its production of sphingolipids, likely as a compensatory mechanism for phospholipid incorporation into the membrane^19^. This adaptation also enhances its resistance to antibiotics and phages. Sphingolipid-deficient *Porphyromonas gingivalis* and *Bacteroides fragilis* which are common in oral and intestinal microbiomes, respectively, have reduced stationary phase survival^20, 21^, and sphingolipid membrane microdomains in *B. fragilis* have been suggested to mediate signal transduction and stress response pathways^21^. Overall, the presence of sphingolipids in bacterial membranes suggests involvement in host-microbe interactions^2, 12, 21, 22^, or generally an adaptation strategy involving changes in the membrane structure or its fluidity. So far, the study of bacterial sphingolipid biosynthesis and function has relied on microbial cultivation. Over 500 headgroup variants and acyl-chain alterations of sphingolipids have been reported from laboratory cultures^23^. However, the lack of cultured representatives of the vast majority of bacteria prevents comprehensive sphingolipid characterization throughout the tree of life, limiting our understanding of the structural diversity of sphingolipids, and of their molecular and ecological function^24^. Bacterial sphingolipids are often associated with eukaryotic hosts and rarely reported in free-living environmental microbiomes, and the extent of structural diversity of bacterial sphingolipids and the identity of producers in the environment remains unknown. The eukaryotic alga *Emiliania huxleyi* is a known contributor to sphingolipid production in marine ecosystems upon viral infection^25^. The virus prompts the synthesis of distinct glycosphingolipids that trigger host-programmed cell death^22, 26, 27^. In hydrothermal marine sediments off the coast of Greece, various types of phosphosphingolipids have been found^28^. And in the deep waters of the Eastern Tropical North Pacific Ocean and in the anoxic and sulfidic (euxinic) zone of the Black Sea, glycosphingolipids of unknown source have been detected^28, 29^.

**SUPPLEMENTARY FIGURES**

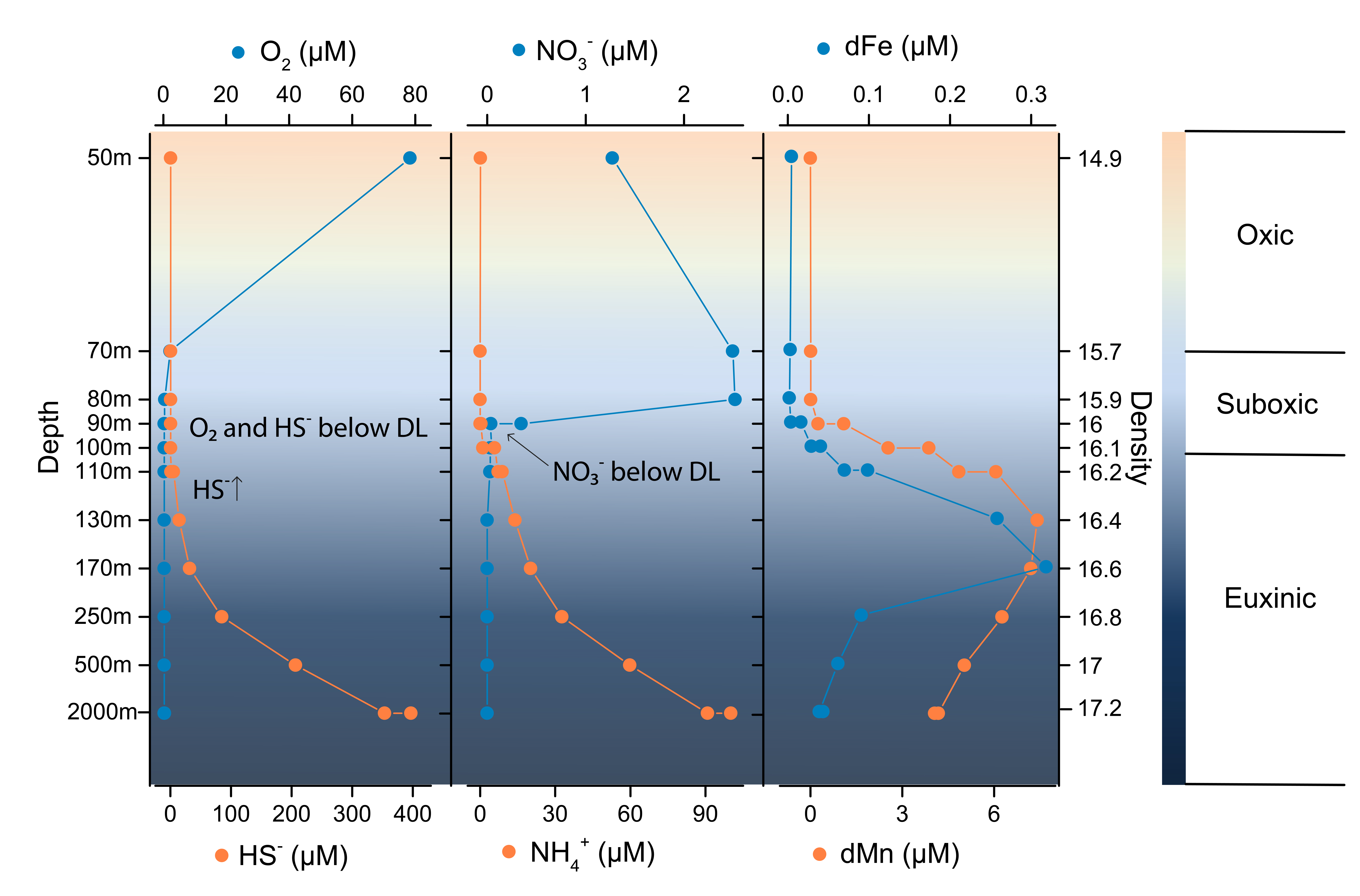

**Supplementary Fig. 1 | Hydrographic and chemical parameters across the water column (depth 50-2000 mbsl) of the Black Sea.** (**a**) Concentrations of $O_{2}$ and $H_{2}S$ (µM). (**b**) Concentrations of $\mathrm{NO}_{3}^{-}$ and $\mathrm{NH}_{4}^{+}$ (µM). (**c**) Concentrations of dissolved Mn (dMn, µM) and Fe (dFe, µM). Data of a and b are from ref. ^30^, data of c were from ref. ^31^.

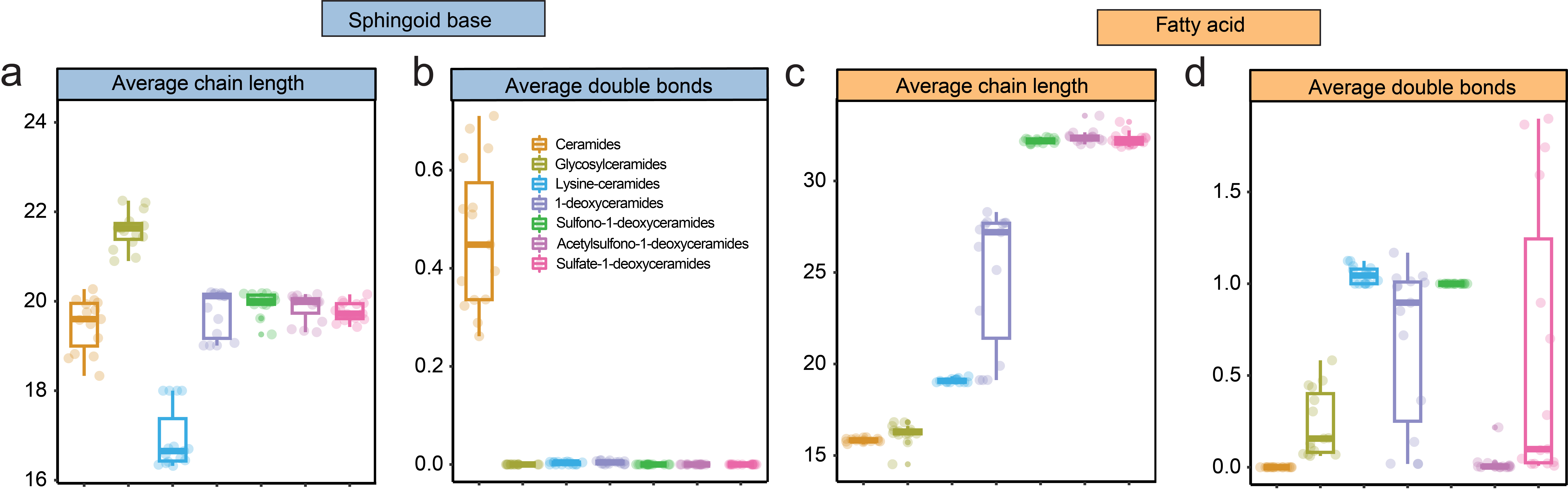

**Supplementary Fig. 2 | Detailed structure of sphingolipid classes based on 15 samples average abundance.** (**a**) The average carbon atoms of the chain length of the sphingoid bases. (**b**) The average number of double bond equivalents (DBEs) of the sphingoid base. (**c**) The average carbon atoms of the chain length of the fatty acid. (**d**) The average number of DBEs of the fatty acid.

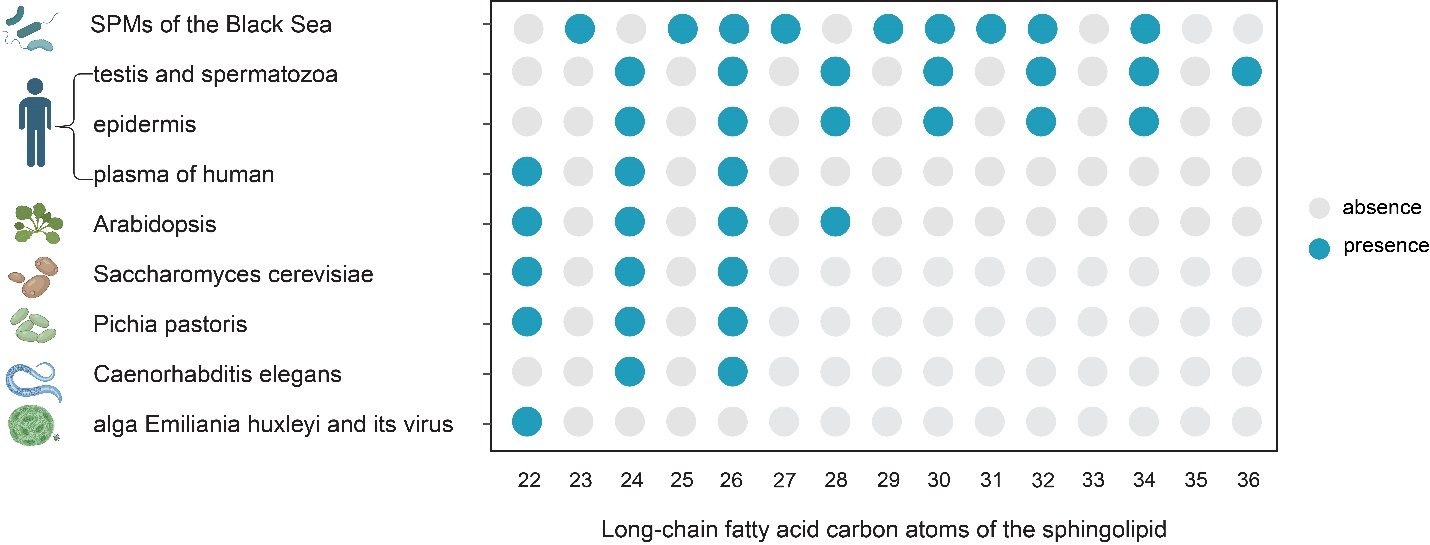

**Supplementary Fig. 3 | Sphingolipids with long-chain fatty acid group (carbon atom** $\boldsymbol{\geq}$ **22) found in the water column of the Black Sea (this study), mammals^5, 32-36^, plants^9, 37-40^, invertebrates^41^, fungi^39, 42-44^, alga *Emiliania huxleyi* and its virus^22, 25, 26^.**

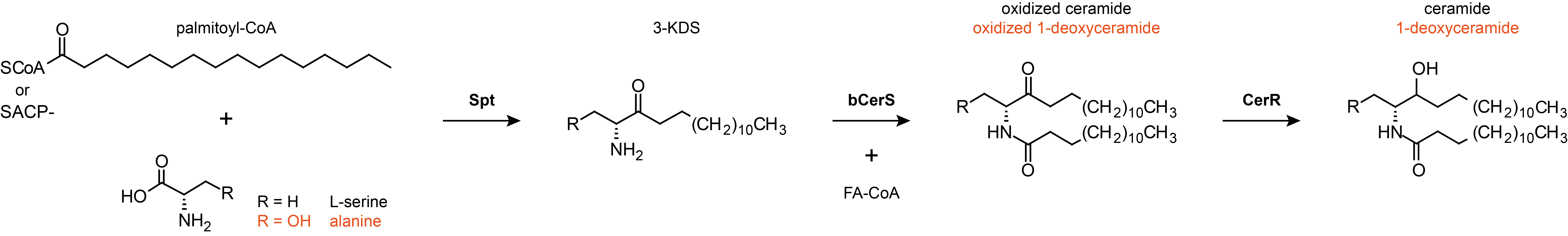

**Supplementary Fig. 4 | The sphingolipid biosynthesis pathway in *Caulobacter crescentus*.** The three proteins that carry out the enzymatic reactions that lead to ceramide are depicted in bold above the arrows. Homologs of Spt are found in bacteria and eukaryotes, bCerS and CerR are bacterial. The proteins of the ceramide biosynthesis pathway were independently identified in ref. ^14^ and ref. ^45^, the names and reaction sequence are from ref. ^14^. 1-deoxyceramides have been shown to be produced by the same set of enzymes when alanine is used as a substrate instead of L-serine^46^. Spt, serine palmitoyltransferase; bCerS, bacterial ceramide synthase; CerR, ceramide reductase.

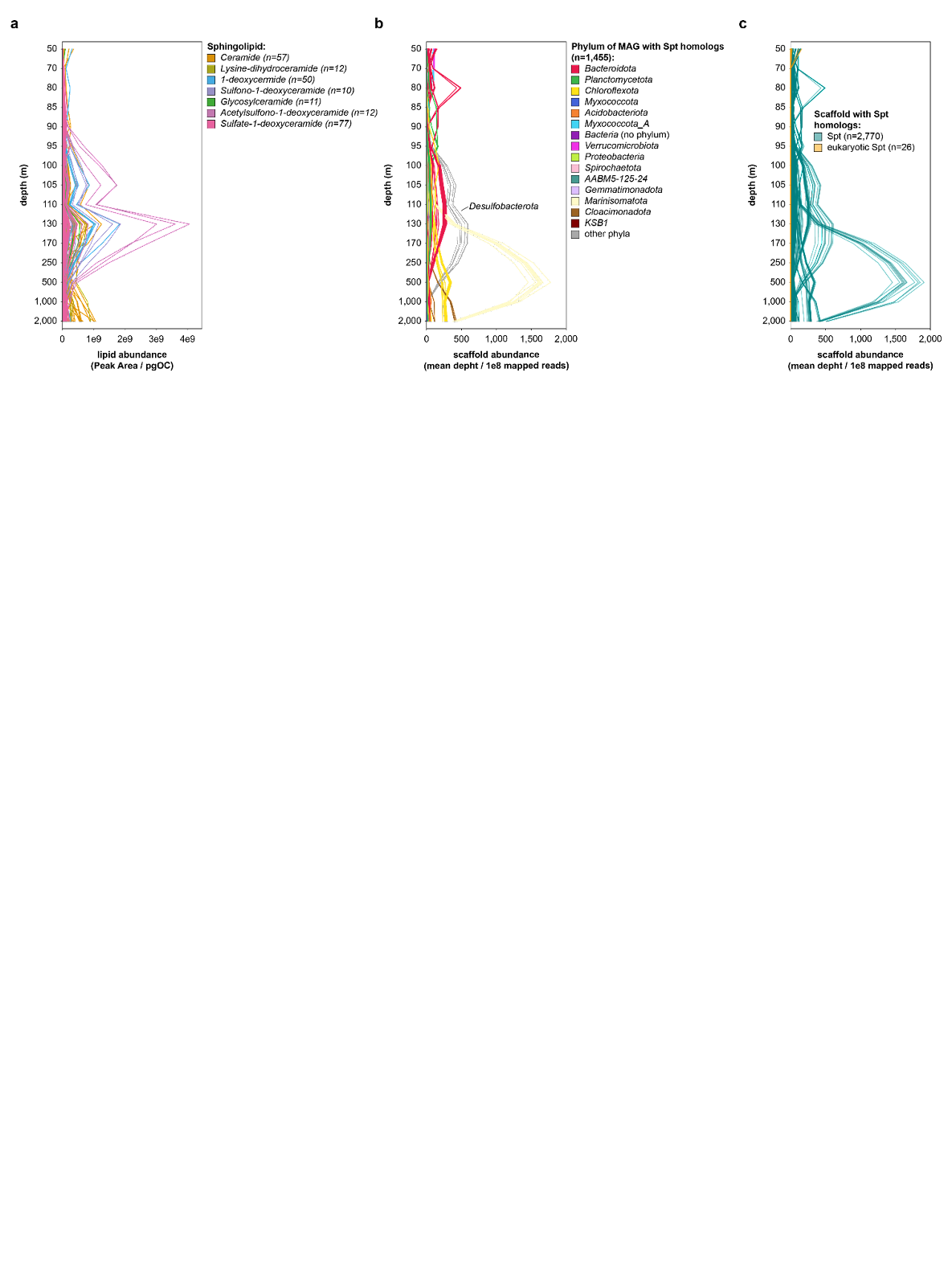

**Supplementary Fig. 5 | Relative abundance profile of sphingolipids and scaffolds containing Spt-like proteins** **throughout the water column.** (**a**) Abundance of sphingolipids. (**b**) Relative abundance of scaffolds that are binned into MAGs. (**c**) Relative abundance of all scaffolds with Spt-like proteins, including those encoding hits to eukaryotic Spt.

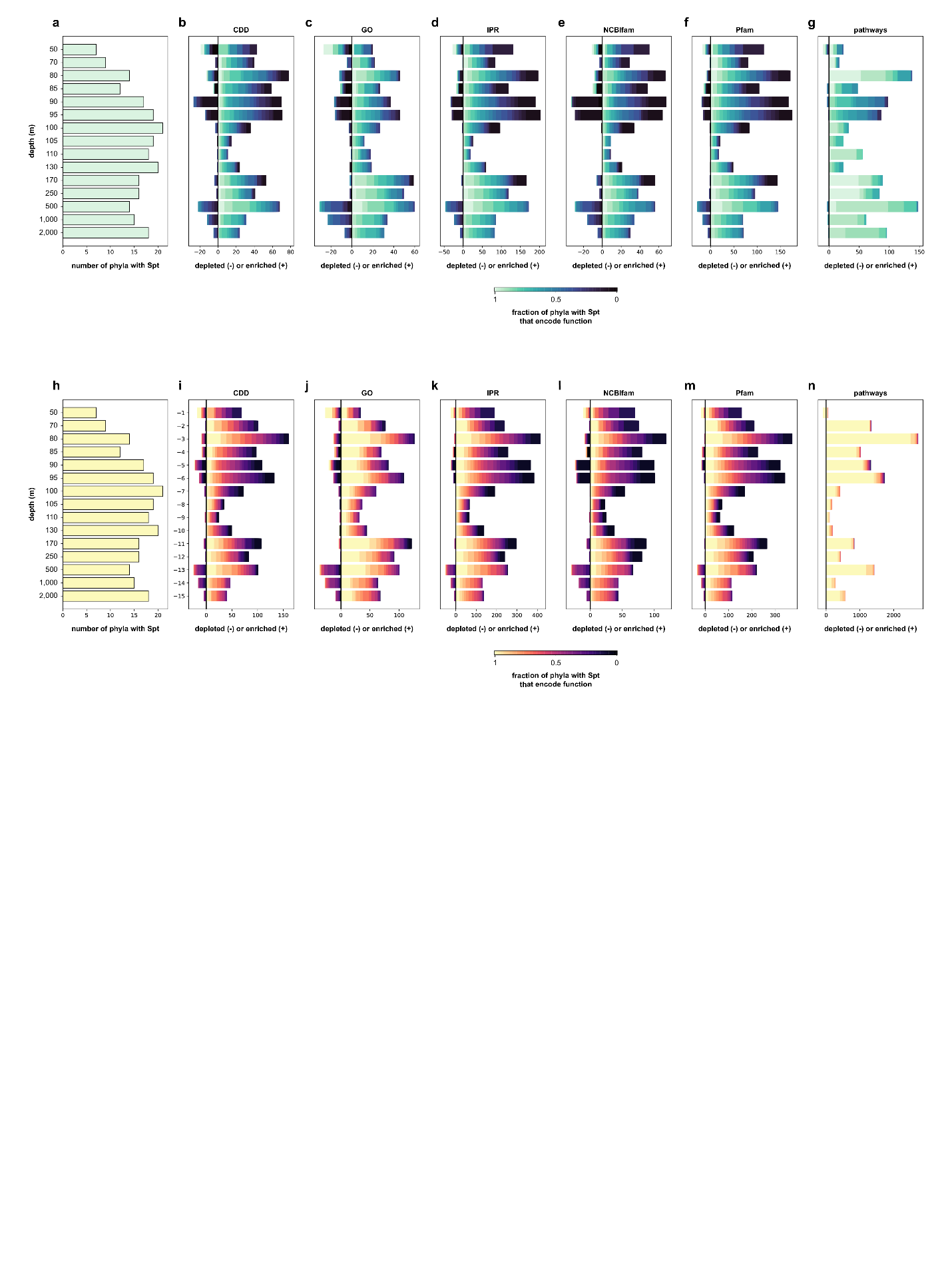

**Supplementary Fig. 6 | Pan-genome-wide association analyses identify functions that are depleted and enriched in MAGs that contain Spt-like proteins.** (**a**) Number of phyla represented by the 704 MAGs that contain Spt-like proteins and whose (completeness – 5 x contamination) ≥ 70%. (**b-g**) Number of functions in different functional universes that are depleted (negative values) or enriched (positive values) in MAGs containing Spt-like proteins compared to MAGs that do not contain Spt-like proteins based on the (qualitative) Fisher’s exact test. The test compares all medium- to high-quality MAGs that were binned from a specific depth. Different panels depict annotations from different functional universes provided by InterProScan: Conserved Domain Database (b), Gene Ontology (c), InterProScan (d), NCBIfam (e), Pfam (f), and pathways (g). Functions are colored according to the fraction of phyla with Spt-like proteins at that depth that contain at least one member that encodes the function. (h) Idem to panel a. (**i-n**) Idem to panels b-g but based on the (quantitative) Mann-Whitney U test. All depleted and enriched functions can be found in Supplementary Data 4.

**SUPPLEMENTARY TABLES**

**Supplementary Table 1 | Lipid standards information used for quantification of major lipid classes, including their full name, molecular formula, and exact mass.** All the standards except GTGT C46^47^ were ordered from Avanti Polar Lipids company.

| **Standard** | **Full name** | **Molecular Formula** | **Exact Mass** |
| --- | --- | --- | --- |
| DGTS-d9 | 1,2-dipalmitoyl-sn-glycero-3-O-4'-[N,N,N-trimethyl(d9)]-homoserine | C_42_H_73_D_9_NO_7_ | 720.657 |
| C16 ceramide(d18:1/16:0) | N-palmitoyl-D-erythro-sphingosine | C_34_H_67_NO_3_ | 537.512 |
| C18 ceramide(d18:1/18:0) | N-stearoyl-D-erythro-sphingosine | C_36_H_71_NO_3_ | 565.543 |
| C24 ceramide(d18:1/24:0) | N-lignoceroyl-D-erythro-sphingosine | C_42_H_83_NO_3_ | 649.637 |
| C24:1 ceramide(d18:1/24:1(15Z)) | N-nervonoyl-D-erythro-sphingosine | C_42_H_81_NO_3_ | 647.622 |
| Glucosyl(β)C12 Ceramide | N-(dodecanoyl)-1-ß-glucosyl-sphing-4-ene | C_36_H_69_NO_8_ | 643.502 |
| Hydrogenated MGDG | Monogalactosyldiacylglycerol (plant hydrogenated) | C_43_H_82_O_10_ | 758.591 |
| Hydrogenated DGDG | Digalactosyldiacylglycerol (plant hydrogenated) | C_51_H_96_O_15_ | 948.675 |
| DGTS (32:0) | 1,2-dipalmitoyl-sn-glycero-3-O-4'-(N,N,N-trimethyl)-homoserine | C_42_H_81_NO_7_ | 711.601 |
| 1-deoxyceramide(d18:1/24:0) | N-[(2S,3R,4E)-3-Hydroxy-4-octadecen-2-yl] tetracosanamide | C_42_H_83_NO_2_ | 633.642 |
| GTGT C46 | C46 glycerol trialkyl glycerol tetraether | C_46_H_94_O_6_ | 742.704 |
| 15:0-18:1(d7) PC | 1-pentadecanoyl-2-oleoyl(d7)-sn-glycero-3-phosphocholine | C_41_H_74_D_7_NO_8_P | 752.606 |
| 15:0-18:1(d7) PE | 1-pentadecanoyl-2-oleoyl(d7)-sn-glycero-3-phosphoethanolamine | C_38_H_67_D_7_NO_8_P | 710.559 |
| 15:0-18:1(d7) PG | 1-pentadecanoyl-2-oleoyl(d7)-sn-glycero-3-[phospho-rac-(1'-glycerol)] (sodiumsalt) | C_39_H_68_D_7_O_10_P | 763.535 |
| 15:0-18:1(d7) PI | 1-pentadecanoyl-2-oleoyl(d7)-sn-glycero-3-phosphoinositol(ammoniumsalt) | C_42_H_72_D_7_O_13_P | 846.596 |
| 18:1(d7) LPC | 1-oleoyl(d7)-2-hydroxy-sn-glycero-3-phosphocholine | C_26_H_45_D_7_NO_7_P | 528.392 |
| 18:1(d7) LPE | 1-oleoyl(d7)-2-hydroxy-sn-glycero-3-phosphoethanolamine | C_23_H_39_D_7_NO_7_P | 486.345 |
| 18:1(d7) MG | 1-oleoyl(d7)-rac-glycerol | C_21_H_33_D_7_O_4_ | 363.336 |
| 15:0-18:1(d7) DAG | 1-pentadecanoyl-2-oleyol(d7)-sn-glycerol | C_36_H_61_D_7_O_5_ | 587.550 |
| 15:0-18:1(d7)-15:0 TAG | 1,3-dipentadecanoyl-2-oleyol(d7)-glycerol | C_51_H_89_D_7_O_6_ | 811.764 |
| 18:1(d9) SM | N-oleoyl(d9)-D-erythro-sphingosylphosphorylcholine | C_41_H_72_D_9_N_2_O_6_P | 737.639 |
| Cholesterol (d7) | cholesterol-d7 | C_27_H_39_OD_7_ | 393.398 |

**Supplementary Table 2 | Lipid standards used in curves for quantification of major lipid classes.** Response factors for each class of lipids were approximated by external calibration with standards. The slope of the linear curve (Peak Area/ngOC) as well as the ratio of that slope to the DGTS-d9 reference standard (Response factor relative to DGTS-d9 curve) is also reported. An average response factor of ceramide mixers was used for ceramide related sphingolipid calibration. For the class of lipids with no standards available (e.g., ornithine lipids), a response factor from another class of amino lipid (DGTS) was used for calculation.

| **Standard** | **Used to quantify** | **Peak area/ngOC** | **Response factor** |
| --- | --- | --- | --- |
| DGTS-d9 |  | 1.58E+09 | 1.00 |
| C16 ceramide (d18:1/16:0) | Treat as ceramide mixers, use average response factor to quantify ceramide related lipids | 1.88E+08 | 0.12 |
| C18 ceramide (d18:1/18:0) |  | 2.01E+08 | 0.13 |
| C24 ceramide (d18:1/24:0) |  | 1.64E+08 | 0.10 |
| C24:1 ceramide (d18:1/24:1 (15Z)) |  | 1.85E+08 | 0.12 |
| Ceramide mixers | Ceramide related sphingolipids |  | 0.12 |
| Glucosyl (β) C12 Ceramide | Glycosylceramides | 1.40E+08 | 0.09 |
| 1-deoxyceramide (d18:1/24:0) | 1-deoxyceramide related sphingolipids | 2.25E+08 | 0.14 |
| Hydrogenated MGDG | Monogalactosyldiacylglycerol (MGDG) | 9.54E+07 | 0.06 |
| Hydrogenated DGDG | Digalactosyldiacylglycerol (DGDG) | 1.66E+07 | 0.01 |
| DGTS (32:0) | DGTS (diacylglyceryl trimethylhomoserines) | 9.43E+08 | 0.60 |
|  | DGCC (diacylglyceryl-3-Ocarboxyhydroxymethylcholine) |  |  |
|  | DGTA (diacylglyceryl hydroxymethyl- trimethyl-β- alanine) |  |  |
| GTGT C46 | GDGT (Glycerol dialkyl glycerol tetraether lipids) | 1.17E+09 | 0.74 |
| 15:0-18:1(d7) PC | Phosphatidylcholine (PC) | 6.77E+08 | 0.43 |
| 15:0-18:1(d7) PE | Phosphatidylethanolamine (PE) | 9.67E+07 | 0.06 |
| 15:0-18:1(d7) PG | Phosphatidylglycerol (PG) | 2.63E+07 | 0.02 |
| 15:0-18:1(d7) PI | Phosphatidylinositol (PI) | 1.97E+07 | 0.01 |
| 18:1(d7) LPC | LysoPC | 7.38E+08 | 0.47 |
| 18:1(d7) LPE | LysoPE | 4.15E+07 | 0.03 |
| 18:1(d7) MG | Monoglycerols (MGs) | 5.56E+07 | 0.04 |
| 15:0-18:1(d7) DAG | Diacylglycerols (DAGs) | 2.22E+08 | 0.14 |
|  | Acyletherglycerols (AEGs) |  |  |
|  | Dietherglycerols (DEGs) |  |  |
| 15:0-18:1(d7)-15:0 TAG | Triacylglycerols (TAGs) | 4.22E+08 | 0.27 |
| 18:1(d9) SM | Sphingomyelin (SM) | 4.42E+08 | 0.28 |
| Cholesterol (d7) | Cholesterol | 1.10E+06 | 0.001 |

**SUPPLEMENTARY DATA CAPTIONS**

**Supplementary Data 1 | Tables with bacterial Spt homolog hits, and eukaryotic Spt homolog hits that were not captured by the bacterial Spt search.** Tables contain the CAT annotation of the scaffold on which the hit was found, the BAT annotation if the scaffold was binned, and the final taxonomic assignment of the hit. Read mappings to both the scaffold and MAG are included, which are the direct mappings of the reads from the sample of which the scaffold was assembled. In addition, the read depth from the all-versus-all mappings are included, representing the abundance profile across the water column.

**Supplementary Data 2 | CD-HIT mapping of Spt homologs, MAFFT alignments, IQ-TREE output files, and iTOL annotation files.** The folder contains all sequences represented by clade A in Fig 3a,b—marked as ‘certain_Spt_seqs’—and all sequences represented by clade B minus clade A in Fig. 3a,b—marked as ‘putative_Spt_seqs’. The file ‘rep2phylum.txt’ contains the mappings of the CD-HIT representative sequence to taxonomic annotation based on the Genome Taxonomy Database of the query sequences.

**Supplementary Data 3 | Source data for Fig. 3c-e.**

**Supplementary Data 4 | Pan-genome-wide association analyses.** The folder contains the list of selected MAGs, results of Fisher’s exact tests and Mann-Whitney U tests, and code to subset and interpret those results.

**REFERENCES**

1.	Geiger, O., Padilla-Gómez, J. & López-Lara, I.M. Bacterial Sphingolipids and Sulfonolipids, in *Biogenesis of Fatty Acids, Lipids and Membranes*. (ed. O. Geiger) 123-137 (Springer International Publishing, Cham; 2019).

2.	Harrison, P.J., Dunn, T. & Campopiano, D.J. Sphingolipid biosynthesis in man and microbes. *Natural Product Reports* **35**, 921-954 (2018).

3.	Nelson, D.L. & Cox, M.M. *Lehninger Principles of Biochemistry. 7th Edition*. (W.H. Freeman, New York; 2017).

4.	Ogretmen, B. Sphingolipid metabolism in cancer signalling and therapy. *Nature Reviews Cancer* **18**, 33-50 (2018).

5.	Hannun, Y.A. & Obeid, L.M. Sphingolipids and their metabolism in physiology and disease. *Nature Reviews Molecular Cell Biology* **19**, 175-191 (2018).

6.	Sigruener, A. *et al.* Glycerophospholipid and sphingolipid species and mortality: The ludwigshafen risk and cardiovascular health (LURIC) study. *PLOS ONE* **9**, e85724 (2014).

7.	Maceyka, M. & Spiegel, S. Sphingolipid metabolites in inflammatory disease. *Nature* **510**, 58-67 (2014).

8.	Resemann, H.C. *et al.* Convergence of sphingolipid desaturation across over 500 million years of plant evolution. *Nature Plants* **7**, 219-232 (2021).

9.	Michaelson, L.V., Napier, J.A., Molino, D. & Faure, J.-D. Plant sphingolipids: Their importance in cellular organization and adaption. *Biochimica et Biophysica Acta (BBA) - Molecular and Cell Biology of Lipids* **1861**, 1329-1335 (2016).

10.	Obeid, L.M., Okamoto, Y. & Mao, C. Yeast sphingolipids: metabolism and biology. *Biochimica et Biophysica Acta (BBA) - Molecular and Cell Biology of Lipids* **1585**, 163-171 (2002).

11.	Jenkins, G.M. *et al.* Involvement of yeast sphingolipids in the heat stress response of Saccharomyces cerevisiae. *Journal of Biological Chemistry* **272**, 32566-32572 (1997).

12.	Heaver, S.L., Johnson, E.L. & Ley, R.E. Sphingolipids in host–microbial interactions. *Current Opinion in Microbiology* **43**, 92-99 (2018).

13.	Olsen, I. & Jantzen, E. Sphingolipids in Bacteria and Fungi. *Anaerobe* **7**, 103-112 (2001).

14.	Stankeviciute, G. *et al.* Convergent evolution of bacterial ceramide synthesis. *Nature Chemical Biology* **18**, 305-312 (2022).

15.	Kawahara, K. *et al.* Chemical structure of glycosphingolipids isolated from Sphingomonas paucimobilis. *FEBS Letters* **292**, 107-110 (1991).

16.	Kawahara, K., Moll, H., Knirel, Y.A., Seydel, U. & Zähringer, U. Structural analysis of two glycosphingolipids from the lipopolysaccharide-lacking bacterium Sphingomonas capsulata. *European Journal of Biochemistry* **267**, 1837-1846 (2000).

17.	Ahrendt, T., Wolff, H. & Bode Helge, B. Neutral and phospholipids of the Myxococcus xanthus lipodome during fruiting body formation and germination. *Applied and Environmental Microbiology* **81**, 6538-6547 (2015).

18.	Kaneshiro, E.S., Hunt, S.M. & Watanabe, Y. Bacteriovorax stolpii proliferation and predation without sphingophosphonolipids. *Biochemical and Biophysical Research Communications* **367**, 21-25 (2008).

19.	Stankeviciute, G., Guan, Z., Goldfine, H. & Klein Eric, A. Caulobacter crescentus adapts to phosphate starvation by synthesizing anionic glycoglycerolipids and a novel glycosphingolipid. *mBio* **10**, e00107-00119 (2019).

20.	Moye, Z.D., Valiuskyte, K., Dewhirst, F.E., Nichols, F.C. & Davey, M.E. Synthesis of sphingolipids impacts survival of Porphyromonas gingivalis and the presentation of surface polysaccharides. *Frontiers in Microbiology* **7** (2016).

21.	An, D., Na, C., Bielawski, J., Hannun, Y.A. & Kasper, D.L. Membrane sphingolipids as essential molecular signals for Bacteroides survival in the intestine. *Proceedings of the National Academy of Sciences* **108**, 4666-4671 (2011).

22.	Schleyer, G. *et al.* In plaque-mass spectrometry imaging of a bloom-forming alga during viral infection reveals a metabolic shift towards odd-chain fatty acid lipids. *Nature Microbiology* **4**, 527-538 (2019).

23.	Fahy, E. *et al.* Update of the LIPID MAPS comprehensive classification system for lipids1. *Journal of Lipid Research* **50**, S9-S14 (2009).

24.	Lewis, W.H., Tahon, G., Geesink, P., Sousa, D.Z. & Ettema, T.J.G. Innovations to culturing the uncultured microbial majority. *Nature Reviews Microbiology* **19**, 225-240 (2021).

25.	Vardi, A. *et al.* Viral glycosphingolipids induce lytic infection and cell death in marine phytoplankton. *Science* **326**, 861-865 (2009).

26.	Schleyer, G. *et al.* Lipid biomarkers for algal resistance to viral infection in the ocean. *Proceedings of the National Academy of Sciences* **120**, e2217121120 (2023).

27.	Ziv, C. *et al.* Viral serine palmitoyltransferase induces metabolic switch in sphingolipid biosynthesis and is required for infection of a marine alga. *Proceedings of the National Academy of Sciences* **113**, E1907-E1916 (2016).

28.	Sollich, M. *et al.* Heat stress dictates microbial lipid composition along a thermal gradient in marine sediments. *Frontiers in Microbiology* **8** (2017).

29.	Schubotz, F., Wakeham, S.G., Lipp, J.S., Fredricks, H.F. & Hinrichs, K.-U. Detection of microbial biomass by intact polar membrane lipid analysis in the water column and surface sediments of the Black Sea. *Environmental Microbiology* **11**, 2720-2734 (2009).

30.	Sollai, M., Villanueva, L., Hopmans, E.C., Reichart, G.-J. & Sinninghe Damsté, J.S. A combined lipidomic and 16S rRNA gene amplicon sequencing approach reveals archaeal sources of intact polar lipids in the stratified Black Sea water column. *Geobiology* **17**, 91-109 (2019).

31.	Dijkstra, N. *et al.* Phosphorus dynamics in and below the redoxcline in the Black Sea and implications for phosphorus burial. *Geochimica et Cosmochimica Acta* **222**, 685-703 (2018).

32.	Sandhoff, R. Very long chain sphingolipids: Tissue expression, function and synthesis. *FEBS Letters* **584**, 1907-1913 (2010).

33.	Furland, N.E., Zanetti, S.R., Oresti, G.M., Maldonado, E.N. & Aveldaño, M.I. Ceramides and sphingomyelins with high proportions of very long-chain polyunsaturated fatty acids in mammalian germ cells. *Journal of Biological Chemistry* **282**, 18141-18150 (2007).

34.	Alecu, I. *et al.* Localization of 1-deoxysphingolipids to mitochondria induces mitochondrial dysfunction. *Journal of Lipid Research* **58**, 42-59 (2017).

35.	Zanetti, S.R., de los Ángeles Monclus, M., Rensetti, D.E., Fornés, M.W. & Aveldaño, M.I. Ceramides with 2-hydroxylated, very long-chain polyenoic fatty acids in rodents: From testis to fertilization-competent spermatozoa. *Biochimie* **92**, 1778-1786 (2010).

36.	Vasireddy, V. *et al.* Loss of functional ELOVL4 depletes very long-chain fatty acids (≥C28) and the unique ω-O-acylceramides in skin leading to neonatal death. *Human Molecular Genetics* **16**, 471-482 (2007).

37.	Luttgeharm, K.D., Kimberlin, A.N. & Cahoon, E.B. Plant sphingolipid metabolism and function, in *Lipids in Plant and Algae Development*. (eds. Y. Nakamura & Y. Li-Beisson) 249-286 (Springer International Publishing, Cham; 2016).

38.	Markham, J.E. *et al.* Sphingolipids containing very-long-chain fatty acids define a secretory pathway for specific polar plasma membrane protein targeting in Arabidopsis  *The Plant Cell* **23**, 2362-2378 (2011).

39.	Ternes, P. *et al.* Disruption of the ceramide synthase LOH1 causes spontaneous cell death in Arabidopsis thaliana. *New Phytologist* **192**, 841-854 (2011).

40.	Roudier, F. *et al.* Very-long-chain fatty acids are involved in polar auxin transport and developmental patterning in Arabidopsis  *The Plant Cell* **22**, 364-375 (2010).

41.	Menuz, V. *et al.* Protection of C. elegans from anoxia by HYL-2 ceramide synthase. *Science* **324**, 381-384 (2009).

42.	Schneiter, R. *et al.* Electrospray ionization tandem mass spectrometry (ESI-MS/MS) analysis of the lipid molecular species composition of yeast subcellular membranes reveals acyl chain-based sorting/remodeling of distinct molecular species en route to the plasma membrane. *Journal of Cell Biology* **146**, 741-754 (1999).

43.	Aguilera-Romero, A., Gehin, C. & Riezman, H. Sphingolipid homeostasis in the web of metabolic routes. *Biochimica et Biophysica Acta (BBA) - Molecular and Cell Biology of Lipids* **1841**, 647-656 (2014).

44.	Guillas, I. *et al.* C26-CoA-dependent ceramide synthesis of Saccharomyces cerevisiae is operated by Lag1p and Lac1p. *EMBO Journal* **20**, 2655-2665 (2001).

45.	Olea-Ozuna, R.J. *et al.* Five structural genes required for ceramide synthesis in Caulobacter and for bacterial survival. *Environmental Microbiology* **23**, 143-159 (2021).

46.	Ryan, E., Gonzalez Pastor, B., Gethings, L.A., Clarke, D.J. & Joyce, S.A. in Metabolites, Vol. 13 (2023).

47.	Huguet, C. *et al.* An improved method to determine the absolute abundance of glycerol dibiphytanyl glycerol tetraether lipids. *Organic Geochemistry* **37**, 1036-1041 (2006).
