## Supplementary Data 4 for "Coupled metalipidomics-metagenomics reveal structurally diverse sphingolipids produced by a wide variety of marine bacteria": explore_Fishers_exact_test_results.py

```
#!/usr/bin/env python3

import argparse
import os
import sys


def parse_arguments():
    parser = argparse.ArgumentParser(
            'explore_Fishers_exact_test_results',
            description='Display Fishers exact test results nicer.',
            usage=(
                './explore_Fishers_exact_test_results.py '
                '-s SAMPLE --enr_or_dep enrichment/depletion '
                '[--only_certain, --db DB, --f_phyla FLOAT, -h / --help]'
                ),
            add_help=False
            )

    required = parser.add_argument_group('Required arguments')
    required.add_argument(
            '-s',
            '--sample',
            dest='sample',
            metavar='',
            required=True,
            type=str,
            help='sample, e.g. BlackSea1'
            )
    required.add_argument(
            '--enr_or_dep',
            dest='enr_or_dep',
            metavar='',
            required=True,
            type=str,
            help='enrichment or depletion'
            )

    optional = parser.add_argument_group('Optional arguments')
    optional.add_argument(
            '--only_certain',
            dest='only_certain',
            required=False,
            action='store_true',
            help='display the only_certain results'
            )
    optional.add_argument(
            '--db',
            dest='db',
            metavar='',
            required=False,
            type=str,
            help=('only show results for this database type '
                '[CDD / GO / IPR / NCBIfam / pathways / Pfam]')
            )
    optional.add_argument(
            '--f_phyla',
            dest='f_phyla',
            metavar='',
            required=False,
            type=float,
            help='cut-off for fraction of Spt phyla with function'
            )
    optional.add_argument(
            '-h',
            '--help',
            action='help',
            help='show this help message and exit'
            )

    args, extra_args = parser.parse_known_args()
    if len(extra_args) != 0:
        sys.exit('error: too many arguments supplied.\n{0}'.format(
            '\n'.join(extra_args)))

    if (
            'BlackSea' not in args.sample or
            args.sample.split('BlackSea')[1] not in [
                str(i) for i in range(1, 16)]
            ):
        sys.exit('error: -s SAMPLE should be BlackSea{1..15}.')
    if args.enr_or_dep != 'enrichment' and args.enr_or_dep != 'depletion':
        sys.exit("error: --enr_or_dep should be enrichment or depletion")
    if args.db:
        if (
                args.db != 'CDD' and
                args.db != 'GO' and
                args.db != 'IPR' and
                args.db != 'NCBIfam' and
                args.db != 'pathways' and
                args.db != 'Pfam'
                ):
            sys.exit('error: --db should be '
                    'CDD or GO or NCBIfam or pathways or Pfam')
    if args.f_phyla and (args.f_phyla > 1 or args.f_phyla < 0):
        sys.exit('error: --f_phyla should be >= 0 and <= 1')
        
    return args


def import_mappings():
    mappings = {}

    dir_ = './InterProScan.restructured/mappings'
    for file_ in os.listdir(dir_):
        if not file_.endswith('.mappings.txt'):
            continue

        with open(f'{dir_}/{file_}', 'r') as f:
            for line in f:
                line = line.rstrip().split('\t')

                id_ = (line[0], line[1])
                if id_ in mappings:
                    sys.exit('Someting wrong!')
                mappings[id_] = line[2:]

    return mappings


def import_Caulobacter_vibrioides_NA1000_annotations():
    id2C_vibrioides_NA1000_ORFs = {}

    file_ = ('./Caulobacter_vibrioides_NA1000/InterProScan/'
            'GCF_000022005.1_ASM2200v1_protein.faa.tsv')
    with open(file_, 'r') as f:
        for line in f:
            line = line.rstrip().split('\t')

            id_ = (line[3], line[4])

            id2C_vibrioides_NA1000_ORFs.setdefault(id_, [])
            id2C_vibrioides_NA1000_ORFs[id_].append(line[0])

    return id2C_vibrioides_NA1000_ORFs


def import_Fishers_exact_test_results(sample, only_certain):
    D = {}

    dir_ = './Fishers_exact_test'
    for file_ in os.listdir(dir_):
        if not file_.startswith(f'sample_{sample}.InterProScan'):
            continue

        if not only_certain and not file_.endswith('.Fishers_exact_test'):
            continue
        if only_certain and not file_.endswith(
                '.Fishers_exact_test.only_certain'):
            continue

        with open(f'{dir_}/{file_}', 'r') as f:
            for n, line in enumerate(f):
                if n == 0:
                    continue

                line = line.rstrip().split('\t')

                id_ = (line[0], line[1])
                if id_ in D:
                    sys.exit('Something wrong! (1)')
                D[id_] = line[2:]

    return D


def print_Fishers_exact_test_results_nicely(
        D,
        db,
        p_value_cutoff,
        enr_or_dep,
        f_phyla,
        mappings,
        id2C_vibrioides_NA1000_ORFs
        ):
    i = 0
    for analysis, signature_accession in sorted(
            D, key=lambda x: float(D[x][9])):
        (
                Spt_and_function,
                noSpt_and_function,
                Spt_and_nofunction,
                noSpt_and_nofunction,
                odds_ratio,
                log_odds_ratio,
                pos_or_neg,
                p_value,
                n,
                corrected_p_value,
                Spt_phyla,
                Spt_and_function_phyla
                ) = D[(analysis, signature_accession)]

        if db and analysis != db:
            continue

        if float(corrected_p_value) >= p_value_cutoff:
            continue

        if pos_or_neg != '+' and pos_or_neg != '-':
            sys.exit('Something wrong! (2)')

        if enr_or_dep == 'enrichment' and pos_or_neg == '-':
            continue
        if enr_or_dep == 'depletion' and pos_or_neg == '+':
            continue

        f_of_Spts_with_function = (
                int(Spt_and_function) /
                (int(Spt_and_function) + int(Spt_and_nofunction))
                )
        f_of_noSpts_with_function = (
                int(noSpt_and_function) /
                (int(noSpt_and_function) + int(noSpt_and_nofunction))
                )

        n_Spt_phyla = len(
                [phylum for phylum in Spt_phyla.split('|') if
                    ';p__' in phylum]
                )
        n_Spt_and_function_phyla = len(
                [phylum for phylum in Spt_and_function_phyla.split('|') if
                    ';p__' in phylum]
                )

        if f_phyla and n_Spt_and_function_phyla / n_Spt_phyla < f_phyla:
            continue

        function_in_C_vibrio = False
        if (analysis, signature_accession) in id2C_vibrioides_NA1000_ORFs:
            function_in_C_vibrio = 'yes'
            """
            function_in_C_vibrio = ';'.join(
                    id2C_vibrioides_NA1000_ORFs[
                        (analysis, signature_accession)]
                    )
                    """
        else:
            function_in_C_vibrio = '-'

        print(
                '{0})'
                '\t||'
                '\t{1:.2f}'
                '\t{2:.2f}'
                '\t||'
                '\t{3:.2f}'
                '\t{4:.2f}'
                '\t{5}'
                '\t{6:.2f}'
                '\t||'
                '\t{7}'
                '\t{8}'
                '\t{9:.2f}'
                '\t||'
                '\t{10}'
                '\t||'
                '\t{11}'
                '\t{12}'
                '\t{13}'.format(
                    i,
                    f_of_Spts_with_function,
                    f_of_noSpts_with_function,
                    float(odds_ratio),
                    float(log_odds_ratio),
                    pos_or_neg,
                    float(corrected_p_value),
                    n_Spt_phyla,
                    n_Spt_and_function_phyla,
                    n_Spt_and_function_phyla / n_Spt_phyla,
                    function_in_C_vibrio,
                    analysis,
                    signature_accession,
                    '\t'.join(mappings[(analysis, signature_accession)])
                    )
                )

        i += 1

    return


def main():
    args = parse_arguments()

    mappings = import_mappings()
    (
            id2C_vibrioides_NA1000_ORFs
            ) = import_Caulobacter_vibrioides_NA1000_annotations()
    D = import_Fishers_exact_test_results(args.sample, args.only_certain)

    p_value_cutoff = 0.05
    print_Fishers_exact_test_results_nicely(
            D,
            args.db,
            p_value_cutoff,
            args.enr_or_dep,
            args.f_phyla,
            mappings,
            id2C_vibrioides_NA1000_ORFs
            )


if __name__ == '__main__':
    main()
```
